# The battle of the sexes in humans is highly polygenic

**DOI:** 10.1101/2024.07.23.604850

**Authors:** Jared M. Cole, Carly B. Scott, Mackenzie M. Johnson, Peter R. Golightly, Jedidiah Carlson, Matthew J. Ming, Arbel Harpak, Mark Kirkpatrick

**Author notes:** Correspondence should be addressed to M.K. and A.H.

## Abstract

Sex-differential selection (SDS), which occurs when the fitness effects of alleles differ between males and females, can have profound impacts on the maintenance of genetic variation, disease risk, and other key aspects of natural populations. Because the sexes mix their autosomal genomes each generation, quantifying SDS is not possible using conventional population genetic approaches. Here, we introduce a novel method that exploits subtle sex differences in haplotype frequencies resulting from SDS acting in the current generation. Using data from 300K individuals in the UK Biobank, we estimate the strength of SDS throughout the genome. While only a handful of loci under SDS are individually significant, we uncover polygenic signals of genome-wide SDS for both viability and fecundity. An interesting life-history tradeoff emerges: alleles that increase viability more in one sex increase fecundity more in the other sex. Lastly, we find evidence of SDS on fecundity acting on alleles affecting arm fat-free mass. Taken together, our findings connect the long-standing evidence of SDS acting on human phenotypes with its impact on the genome.

**Significance statement:** Selection often acts differently on females and males, as evidenced by the striking sexual dimorphism found in many taxa. As a result, alleles can have different fitness effects in each sex. Consequences can include higher levels of genetic variation and higher disease burdens in populations. This study introduces a novel method to quantify this sex-differential selection (SDS) and reveals that it acts throughout the human genome. We discovered a life history tradeoff between survival and fecundity in females and males and that SDS on fecundity acts on alleles affecting arm fat-free mass.

## 1 Introduction

Selection that acts differently on the sexes plays central roles in diverse evolutionary processes. Paramount among these is the evolution of sexual dimorphism. The phenotypic differences between females and males can be profound, and in many cases are greater than differences between species for individuals of the same sex (1). Sex-differential selection, or SDS, can maintain genetic variation (2, 3), promote genetic diseases (reviewed in (4)), and drive the origin and subsequent evolution of sex chromosomes (5, 6).

Several lines of evidence suggest that SDS is common. Meta-analyses of selection acting on 89 traits in 34 nonhuman animals estimate that perhaps 20% of quantitative traits experience ongoing antagonistic SDS (7). In humans, SDS has been found on numerous phenotypes such as height, body mass, blood pressure, cholesterol, and age at first birth (8–10). Quantitative traits such as these typically show high genetic correlations between the sexes (11–14), which implies that many alleles contributing to their variation have concordant effects in females and males (although not necessarily the same magnitude of effects; Zhu *et al*. (15)). SDS acting on these phenotypes may therefore favor alternate alleles in females and males (16–18). Further, SDS is not limited to quantitative phenotypes: single genes with major fitness effects that differ between the sexes have been found in *Drosophila* (19–21), salmon (22), cichlid fishes (23), sheep (24), voles (25), and humans (4).

Given this plethora of evidence for SDS, it seems inevitable that it acts on numerous sites across the genome. Confirmation of this simple prediction, however, has proven challenging. A major barrier to studying SDS is that the standard tools of molecular evolution used to detect natural selection are of no use. They are based on patterns of genetic variation that take many generations to accumulate (26). Those methods are unusable for detecting SDS because Mendelian segregation erases any sex differences on autosomes at the start of each generation.

This impasse can be surmounted by studying selection “in real time”: searching for the minute sex differences in allele frequencies generated by SDS acting in the current generation. Using this approach, Lucotte *et al*. (27) estimated allele frequency differences between the sexes throughout the genome and reported larger differences on X chromosomes than autosomes. Cheng and Kirkpatrick (28) detected SDS in the 1000 Genomes dataset (29) by aggregating the signal across the genome. Their methods produced similar results in pipefish (30), guppies (31), rockfish (32), and flycatchers (33).

These findings, however, have not been without controversy. Bissegger *et al*. (34) discovered an important bioinformatic artifact that can produce spurious signals of SDS in the genome. They showed that apparent sex differences in allele frequencies at autosomal SNPs in stickleback fishes are in fact the result of sequencing reads from the sex chromosomes being mis-mapped to the autosomes. This problem is particularly acute in species that do not have assembled Y or W chromosomes in their reference genomes because reads from those sex chromosomes are inevitably mapped to the autosomes. The Bissegger study inspired Mank et al. (35) to suggest that the previous reports of SDS described above were erroneous. In a series of papers, Kasimatis *et al*. (36–38) reviewed the earlier work and searched for sex differences in allele frequencies in two large biobanks. They found no individual SNPs were significant at a genomewide level. Based on that finding and their reinterpretation of previous studies, Kasimatis *et al*. (38) concluded that “we see no evidence of [SDS] generating substantial autosomal allelic divergence between the sexes.” Kasimatis *et al.* (37) concurred with Mank *et al*. (35) in objecting to another aspect of the earlier reports: if the observed sex differences in allele frequencies in adults did result from SDS, the total mortality incurred from viability selection would be overwhelming and demographically unsustainable. Most (or perhaps all) of the allele frequency differences between the sexes, therefore, must be caused by sampling. Cheng and Kirkpatrick (39) disputed many of the conclusions drawn by Mank, Kasimatis, and their colleagues.

More recent studies looking for signals of genome-wide SDS have engaged with these criticisms directly. In an important advance, Ruzicka *et al*. (40) addressed various technical artifacts identified by Mank, Kasimatis, and colleagues and showed that there are significant sex differences in allele frequencies resulting from both viability and fecundity selection in the UK Biobank (41). Once again, while the differences were so small that no individual site in the genome reached statistical significance, SDS was detected by summing the signal across the genome. Ruzicka *et al* also found that certain genomic compartments (e.g. coding regions) are enriched for single nucleotide polymorphisms (SNPs) showing evidence of SDS. Lucotte *et al*. (42), relying on a trio dataset to detect sex-biased transmission distortion from parents to offspring, found candidate regions of SDS primarily located in genes involved in embryonic development. Zhu *et al*. (15) showed that the sex differences in allelic effects on circulating testosterone level correlate with sex differences in allele frequencies, providing a direct link between SDS acting on that phenotype with SDS acting on its underlying loci. Wang *et al*. (43) and Chen *et al*. (44) reported further evidence that sex differences in allele frequencies are enriched on X chromosomes relative to autosomes. Most recently, Chakrabarty *et al*. (45) used a different methodology to report that testosterone drives sexually antagonistic selection on several anthropomorphic traits.

In sum, there seems to be no argument that many phenotypes experience SDS and that this selection must result in genetic differences between the sexes within each generation. Further, compelling evidence now exists for genome-wide signals of SDS. However, we do not yet have an understanding of the genomewide strength of SDS nor the extent of the mortality generated by such selection.

Here we introduce a novel likelihood-based method that leverages information from linked SNPs that have been phased. Phased data, where alleles have been resolved onto maternal chromosomes, allows us to infer linkage disequilibrium between SNPs and use haplotypes in our analysis. In a departure from previous studies (e.g. (27, 28, 40)), rather than focus on descriptive statistics such as *F*_ST_ between the sexes, we estimate the generative parameters of SDS, including selection coefficients and the prevalence of SDS genome-wide. When aggregated across autosomal sites, both viability and fecundity show highly significant signals of SDS, in agreement with Ruzicka *et al* (40). We develop a model that links sex differences in allelic effects to SDS and find evidence that alleles that affect fat-free arm mass are under SDS for fecundity. An intriguing result is that SDS involves a life history tradeoff: alleles that more strongly increase viability in females than males also more strongly increase fecundity in males than in females. We estimate that 20% of autosomal sequence is linked to targets of SDS with selection coefficients on the order of *s* = 10*^−^*^3^, and discuss the implications for the mortality load as a result of SDS.

## 2 Results

Detecting the subtle signals of selection requires large samples. We therefore turned to the UK Biobank, which is the largest available database of whole genome sequences (41). The database relies on active enrollment with participants that tend to be older and healthier than the general population (46). As a result, allele frequency differences between males and females can result from sex differences in participation in the database rather than SDS (47), an issue we return to in the Discussion. We take additional steps to mitigate several statistical artifacts that were identified by Mank, Kasimatis, and colleagues as leading to spurious sex differences in allele frequencies (see SI section A for details). After filtering, our dataset consisted of 554,944 phased genotype array SNPs, sampled from 327,918 female haplotypes and 279,730 male haplotypes (see Methods).

### 2.1 The strength of SDS

Our approach comprises two elements that are novel to studies of SDS. Here we outline them; details are given in the Methods and SI.

The first element is a population genetic model for how SDS with a given selection coefficient drives sex differences in haplotype frequencies (Figure 1A,B). At conception, autosomal allele frequencies are expected to be equal in females and males. SDS acting on a site then generates an allele frequency difference between the sexes at that site, and also at neighboring sites that are in linkage disequilibrium with it. This is a form of genetic hitchhiking (48) that occurs within the current generation.

**Figure 1:**
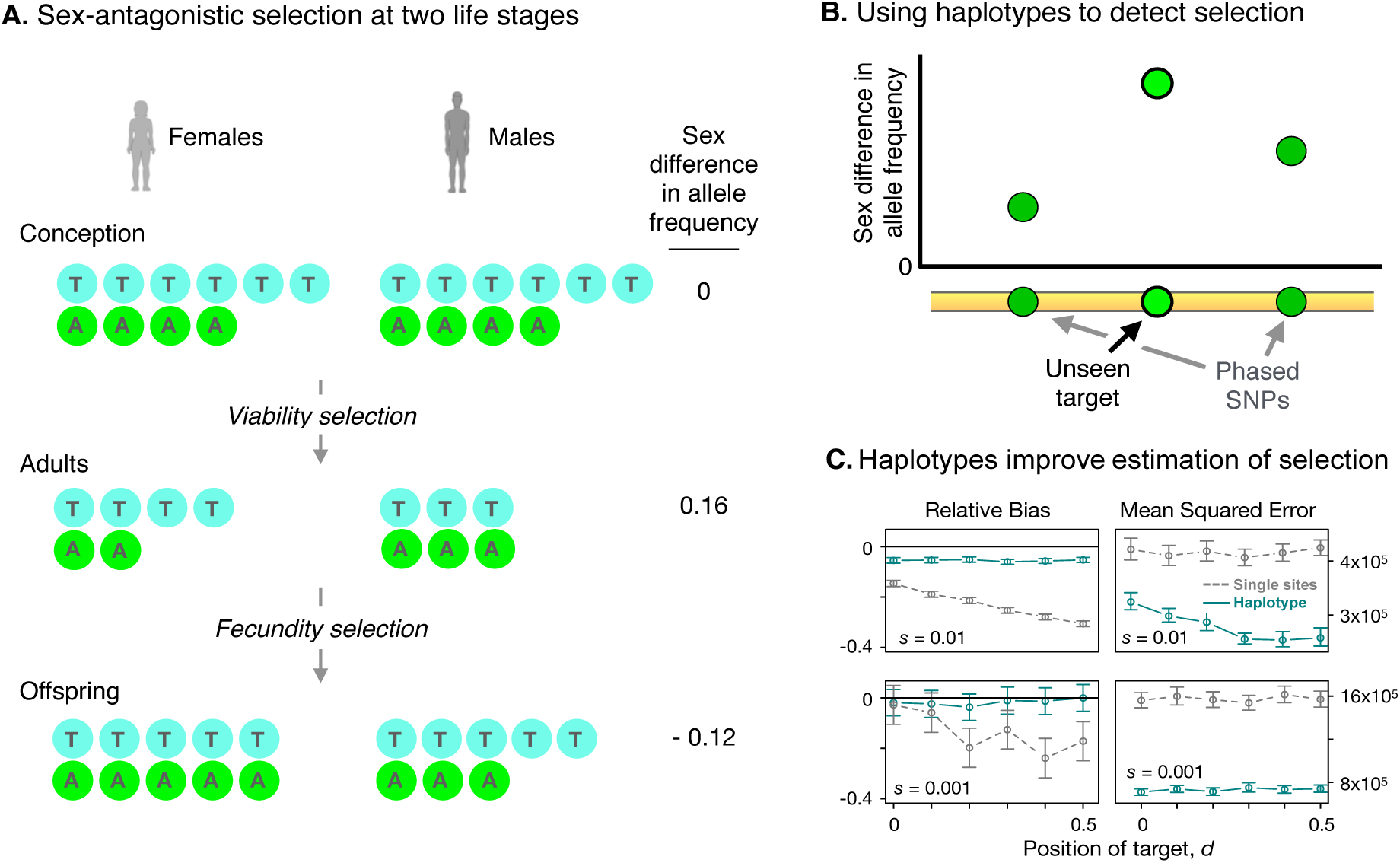
Haplotype-based estimation of SDS across two life stages. **(A)** Alleles T and A are equal in frequency in males and females at conception. Sex-differential viability selection generates allele frequency differences between sexes in the adults. In this example, the T allele is favored in females by viability selection and in males under fecundity selection. **(B)** Using phased SNPs allows estimation of selection acting at unobserved sites. SDS at the unseen target site causes allele frequencies to differ between the sexes at that site and at observed sites in LD with it. **(C)** Simulations show the haplotype method has improved performance relative to single-site approaches. The left-hand panels show the relative bias (proportional deviation from the true selection coefficient) of the haplotype vs. single site approaches; the right-hand panels show the mean squared error. The horizontal axis shows the location of the true target of selection. The haplotype approach assumes the unseen target lies midway between the two observed flanking sites (*d* = 0.5). The single site approach assumes the target is one of the observed sites (here assumed to be the right hand site: *d* = 0).

We reasoned that very few of the targets of SDS would be among the phased SNPs we studied because they represent less than 1% of all SNPs. We therefore assumed targets of SDS lie between observed pairs of phased SNPs. Our model estimates the strength of SDS acting on these unseen targets based on the sex differences in allele frequencies at the observed pair of flanking SNPs and the linkage disequilibrium between those two sites. Simulations show that the resulting estimates are robust to violations of our assumption about the target’s location (Figure 1C; SI section A.3).

The selection coefficients reflecting viability selection acting on females and males could be estimated from changes in allele frequencies between conception and adulthood. We do not know the frequencies at conception, however, so we assume that SDS on viability is antagonistic and sex-symmetric; that is, the selection coefficients in the two sexes are equal in magnitude but opposite in sign. In that case, the frequencies at conception are the average frequencies across female and male adults. SDS that is not sex-symmetric will bias the estimates, but the relative bias is expected to be very small (SI section A.3). Heterozygotes are assumed to have intermediate fitness, and if this is not the case then the selection coefficients represent the average fitness effects. Finally, the model assumes there is no epistasis. Under these conditions, basic population genetics theory predicts the frequencies in surviving adults of the haplotypes consisting of three SNPs (the unseen target and two observed flanking SNPs), given the viability selection coefficient.

This population genetic model is teamed up with the second element of our approach: a likelihood model to estimate the selection coefficients. Assuming that individuals in the UK Biobank represent a random sample from the population, the likelihood of observing given numbers of each haplotype is given by the multinomial distribution, based on the frequencies predicted by the population genetic model. We estimated the selection coefficients by maximizing this likelihood.

A similar strategy is used to estimate the strengths of fecundity selection and total selection over the lifetime. The frequencies of haplotypes in the sample of adults are weighted by the numbers of children individuals reported. These weighted frequencies are used to estimate selection coefficients pertaining to total selection. The fecundity selection coefficients are then calculated as the difference between the total and viability selection coefficients. At all three life stages (conception, adults, offspring), we assume the population is sufficiently large that random deviations from the expected haplotype frequencies (that is, drift occurring in a single generation) can be ignored. This assumption is plausible: with a population size of 450,000 (the approximate number of newborns identified as white, British between 2007-2010 (49) and a minor allele frequency of 0.05, the Wright-Fisher binomial sampling variance is only 10*^−^*^7^ (corresponding to a coefficient of variation of 0.006).

We used simulations to evaluate how our haplotype approach performs relative to one that uses only allele frequencies at single sites. We simulated 3-site haplotypes with known selection coefficients under varying values for the minor allele frequencies and linkage disequilibrium drawn from the UK Biobank (see Methods), with the unseen target of selection at different locations between the flanking SNPs. We find that our estimator of the selection coefficient is conservative (slightly biased downward) and is robust to violations of the assumptions about the target location. Importantly, it has improved performance relative to those based on single sites: it has a 21% lower mean-squared error and 77% lower relative bias (Figure 1C and SI section B.1).

### 2.2 Signals of SDS are highly polygenic

Viability, fecundity, and total selection all show significant signals of sex-differential selection (Figure 2). Selection coefficients for viability and lifetime reproductive success are significantly enriched for large values (compared to their null distributions). Selection coefficients for all three modes of selection are significantly higher than expected by their respective empirical nulls (Mann-Whitney *p <* 0.001; SI figures B.3.1-B.3.3). Out of 248,059 windows, fifteen reach significance for viability selection after correction (non-sequential Bonferroni, *α* = 0.05, see SI section A.6). Within these windows there are 29 SNPs that fall in or near genes involved in cancer, height, neurodevelopmental and neuropsychiatric disorders, and other functions (SI table B.4.1). Although this number is modest, it improves on previous studies based on single sites, which found no genomic targets of SDS that were individually significant. No sites reach significance for fecundity selection, however (SI section B.3). These findings confirm the previous report that signals of SDS in the UK Biobank are subtle and highly polygenic (40).

**Figure 2:**
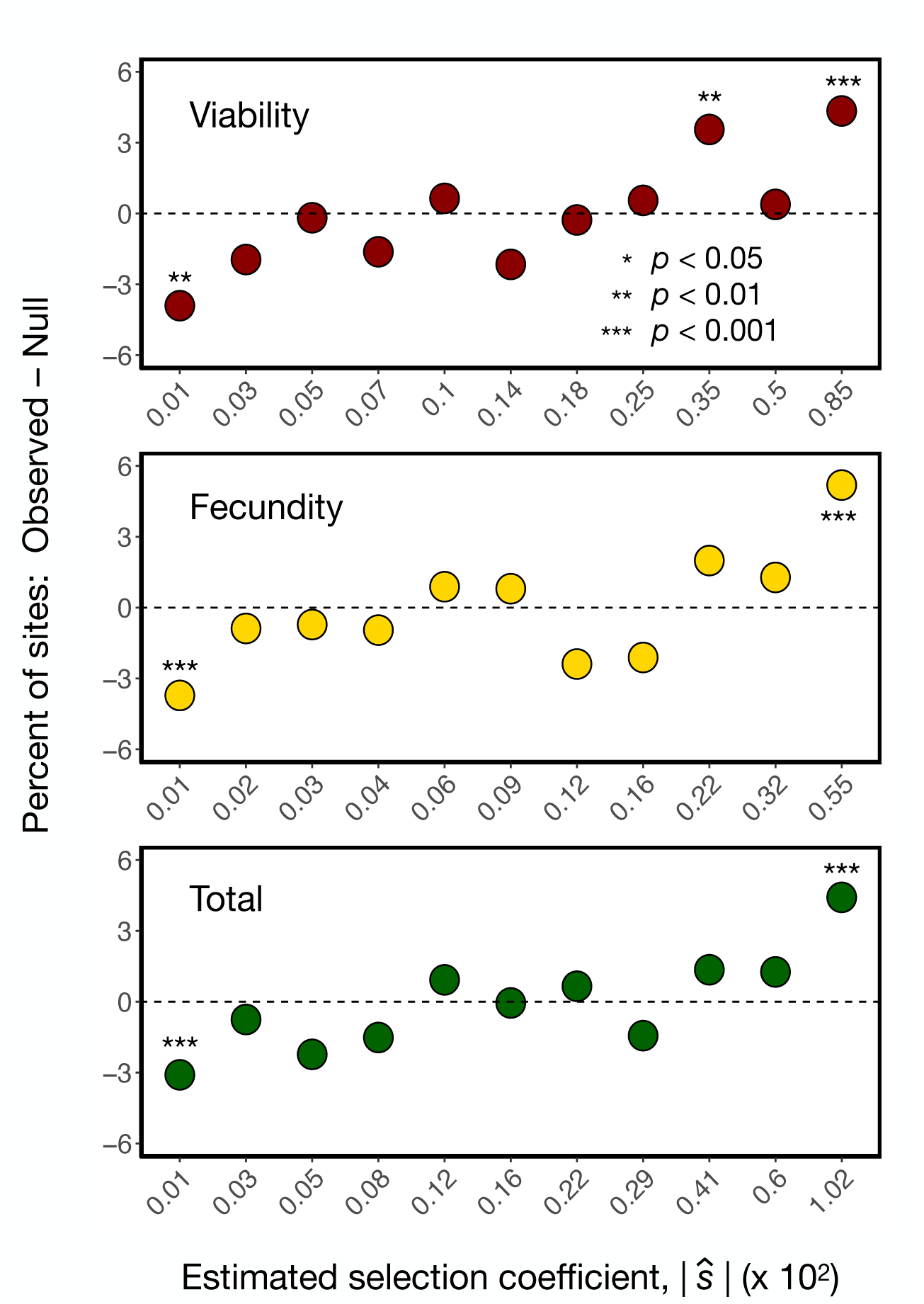
Genome-wide signals of polygenic sex-differential selection. The absolute values of the selection coefficients (*|ŝ*|) estimated from the empirical null distributions are placed into 11 quantiles for each mode of selection. X-axis values show the bin medians (multiplied by 10^2^). The empirical null distributions were generated by permuting the sex labels among haplotypes. The Y-axis indicates the percentage excess or deficit of *|ŝ*| values in each bin from the observed data. The *p* values for each bin were calculated using *χ*^2^ tests.

SDS includes antagonistic selection, acting in opposite directions in females and males, and concordant selection, which acts in the same direction in both sexes but with different strengths. We cannot distinguish between these two types of SDS acting on viability because we do not know the allele frequencies before selection. We do know them, however, when fecundity selection acts. We can exploit that information by analyzing selection on single sites (rather than on haplotypes, as described above). We therefore modified the likelihood model to estimate the fecundity selection coefficients acting on individual sites separately for females and males (see SI section A.5). Relative to the null expectation under no SDS, sexually antagonistic selection will generate an excess of sites with different signs in females and males. Indeed, among sites with the strongest evidence of antagonistic SDS (specifically, the top 1% of sites among the negative values of 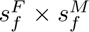, the product of the fecundity selection coefficients for females and males respectively), we find a slight but significant excess (7%; *p <* 0.05 by *χ*^2^) of sites in the observed data compared to the empirical null. This result agrees with Ruzicka *et al*. (40), who found evidence for antagonistic effects using unphased SNPs.

An intriguing discovery is that SDS entails a life-history tradeoff (Figure 3). Selection on alleles that increase survival more strongly in one sex than the other also tend to increase fecundity more strongly in the other sex. Selection coefficients for viability and fecundity are negatively correlated (Pearson *r*, *p* = 0.045), as are the *Z* -scores for the sex differences in selection coefficients obtained by bootstrapping (*p <* 0.001, Methods).

**Figure 3:**
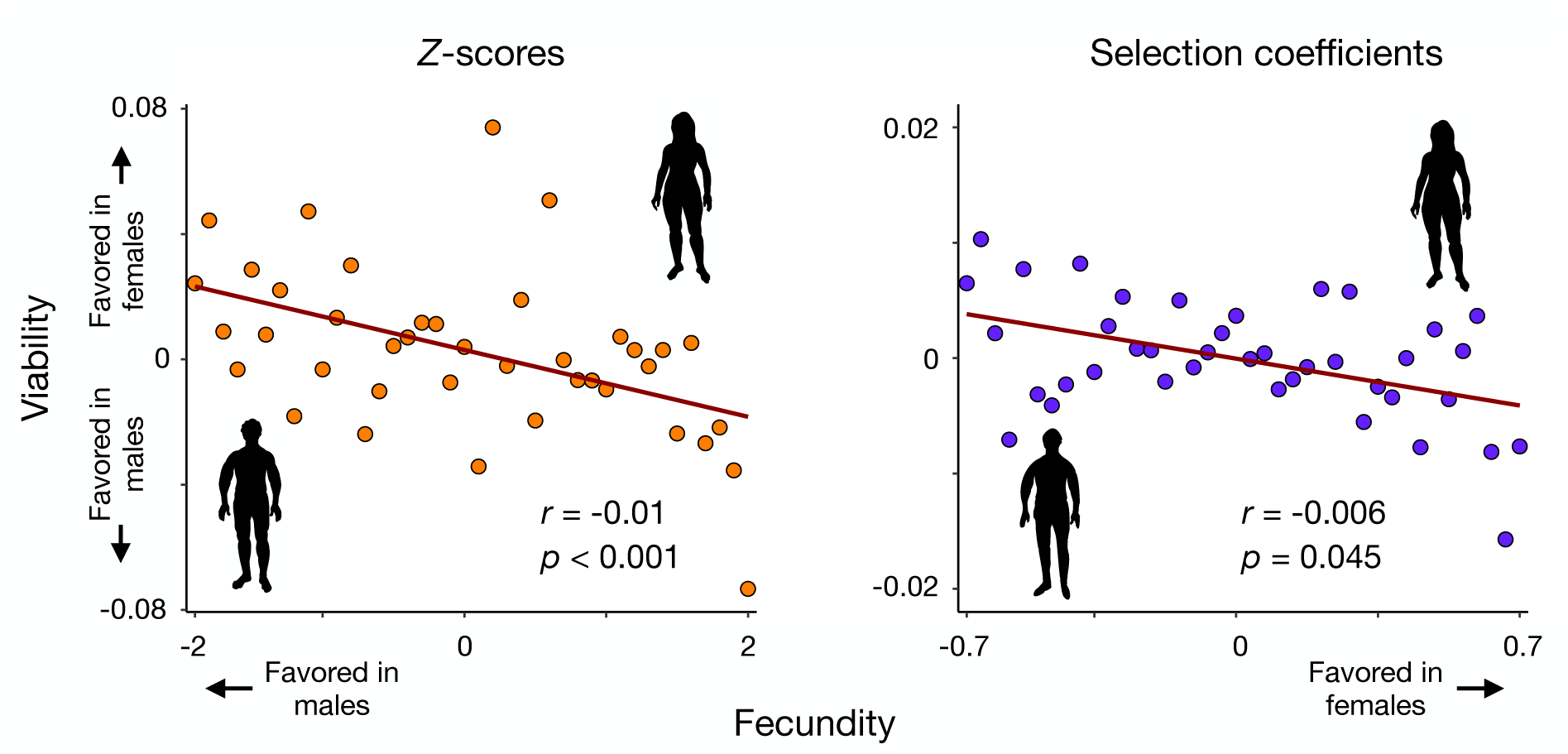
The tradeoff between viability and fecundity selection is sexually antagonistic: alleles that more strongly increase fecundity in females tend to more strongly increase viability in males, and vice versa. Left: The *Z* -scores for viability selection are plotted against the *Z* -scores for fecundity selection. Values are aggregated into 40 bins. Right: The corresponding plot for selection coefficients (multiplied by 10^2^).

Sexually antagonistic selection (SAS) acting on phenotypes generates sex-differential selection on SNPs that contribute to their variation (Figure 4). To maximize our power to detect links between selection acting on phenotypes and genomic sites, we focused on 27 physiological and morphological traits with high SNP heritabilities (15). We leveraged the finding that the human genome is structured into “haplotype blocks” within which linkage disequilibrium is high (50, 51). Therefore to obtain independent estimates, we sampled one pair of phased SNPs from each of the haplotype blocks identified in the European panel of the 1000 Genomes Project by Berisa and Pickrell (52) (see Methods). We modeled the relationship between sex difference of a SNP’s effects on a phenotype and the strength of SDS acting on that SNP (see Methods and SI section A.7). For viability selection, no trait reached statistical significance (*|Z| >* 1.96). The largest signals are for pulse rate (*Z* = 0.54), where alleles with larger effects are favored in females, and testosterone (*Z* = *−*0.69), where alleles with larger effects are favored in males. Signals are stronger for fecundity selection, where selection tends to favor larger effects in males for traits related to body mass. One trait, arm fat-free mass, reaches statistical significance (*Z* = *−*2.22 for the right arm and *Z* = *−*2.01 for the left arm), though no traits remain significant after FDR correction (see SI section A.7).

**Figure 4:**
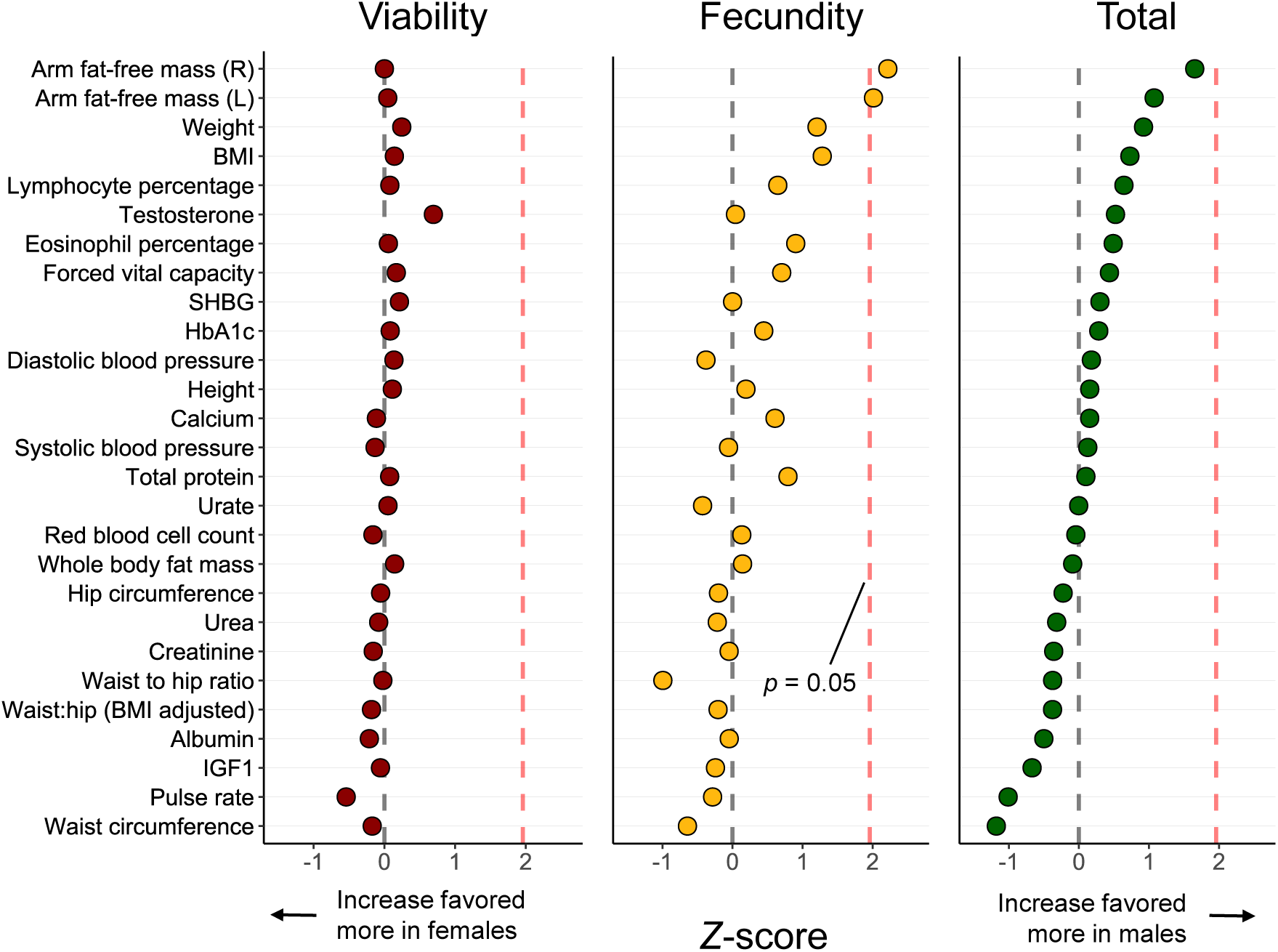
We developed a population genetic model linking sex-specific allelic effects to SDS. The x-axis shows the *Z* -score for the intensity of SDS on alleles affecting the trait. Positive *Z* -scores indicate selection favoring larger (more positive) effects in males, while negative *Z* -scores indicate the opposite.

### 2.3 The strength of SDS, its frequency in the genome, and the mortality it incurs

So far, we have ignored sampling noise, which contributes to sex differences in sample haplotype frequencies—beyond those driven by SDS (28, 35, 37). We therefore used Approximate Bayesian Computation to estimate the average strength of SDS and its frequency in the genome while accounting for sampling noise (See Methods and SI section A.8). We estimate a typical selection coefficient of *s* = 5 *×* 10*^−^*^4^ (90% credible interval [1 *×* 10*^−^*^5^, 3 *×* 10*^−^*^2^]) acting on 20% (90% credible interval [0.01%, 34%]) of haplotype blocks in autosomes (Figure 5A). These results could be used to estimate the mortality load given the allele frequencies at the targets of selection, but these are difficult to estimate (SI section A.8). For example, if a typical minor allele frequency at a site under SDS is 0.01, the mortality load is 0.2% (Figure 5B). The implication is that SDS is widespread across the genome, but may still result in modest mortality.

**Figure 5:**
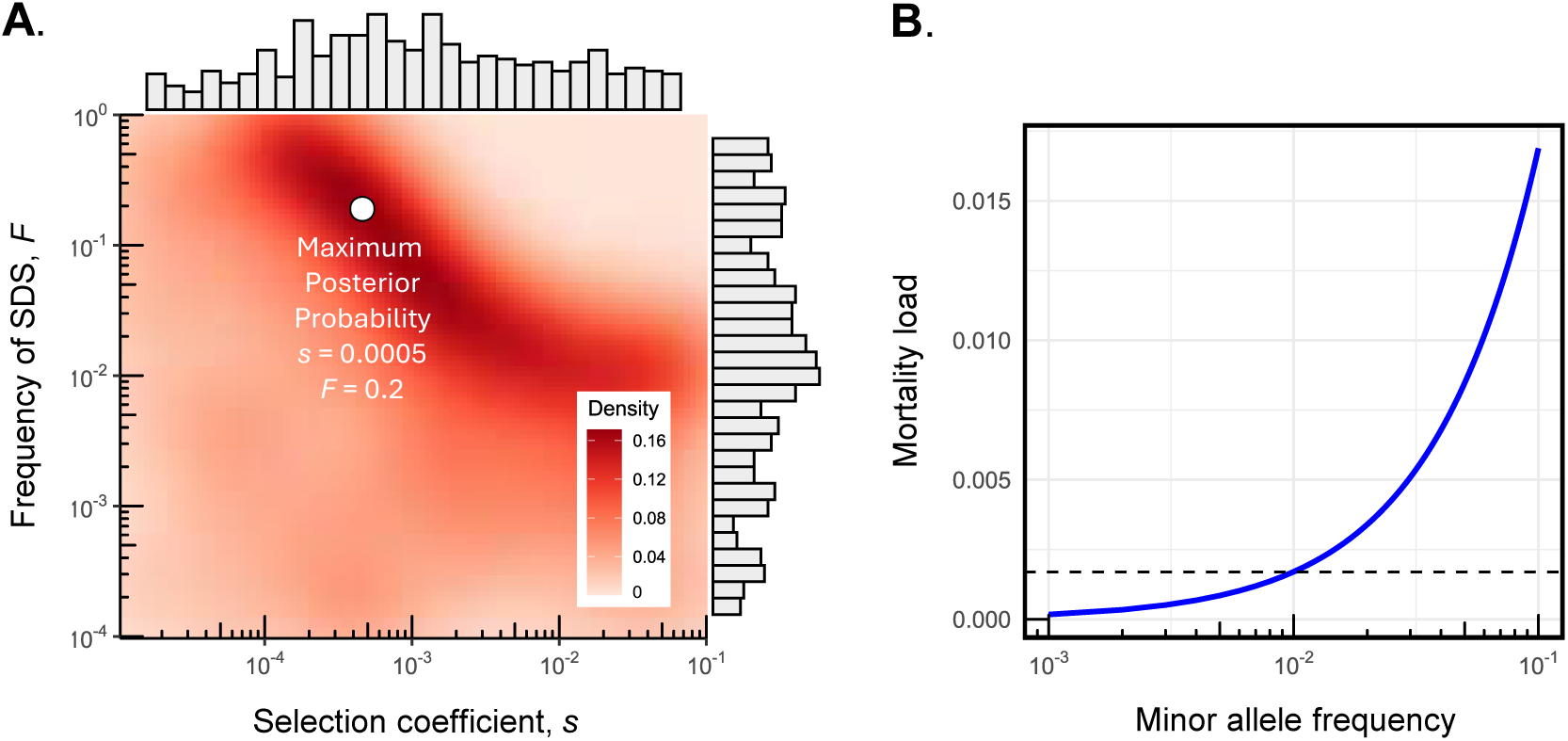
The strength and frequency of SDS and accompanying mortality load. **(A)** Approximate posterior distribution of the frequency of sex-differential selection (*F*) and the selection coefficient for SDS (*s*). F is the probability that a randomly chosen autosomal site is in linkage disequilibrium with a site under SDS. Approximate Bayesian Computation (ABC) allows us to distinguish between allele frequency differences between the sexes caused by SDS and those resulting from sampling. **(B)** The mortality load of SDS given the parameters estimated for *s* and *F* from ABC (panel A) as a function of minor allele frequency at the target of selection. The calculation assumes there are 1,703 independent linkage blocks in the genome. The dotted shows the mortality load (0.2%) when the minor allele frequency at selected sites is 1%.

## 3 Discussion

We find evidence of contemporary sex-differential selection (SDS) across the human genome. Building on previous studies that used intersexual *F*_ST_ to detect SDS (15, 27, 28, 30–33, 40), we developed a population genetic model to estimate the strength of SDS acting on sites within some 550,000 autosomal windows defined by adjacent pairs of phased SNPs. Aggregating these estimates reveals that the cumulative, polygenic effects of SDS on viability and fecundity are significant. SDS may typically generate selection coefficients on the order of *s* = 10*^−^*^3^, with some 20% of autosomal sequence linked to a site under SDS. Of the 27 phenotypes we examined, alleles affecting one (arm fat-free mass) show a significant signal of SDS.

The classic life history tradeoff between viability and fertility has a sexually antagonistic dimension: alleles that increase survival more strongly in one sex tend to increase fecundity more strongly in the other sex. This observation extends an earlier report of a sex-independent tradeoff between viability and fecundity in the UK Biobank (53). Finally, we find that SDS on viability does not necessarily entail a heavy mortality load. The emerging picture is that SDS acts on the genome like “dark energy”: it is ubiquitous but very difficult to observe directly.

Three important caveats pertain to our conclusions. The sex differences in adult haplotype frequencies that are the focus of our method may result from population structure or sex biases in recruitment (47, 54, 55). We have attempted to adjust for population structure statistically (see Methods). Furthermore, a recent analysis found no significant interactions with sex in terms of genetic associations related to UK Biobank participation (56). Nonetheless, our conclusions would be greatly strengthened by replication across other datasets. A second issue involves our inferences regarding phenotypic targets of sex-differential selection. As with all other studies based on correlations between phenotypes and fitness, we do not know if the targets of selection are the traits we are studying or unseen traits that are genetically correlated with them.

A third caveat is that our mortality load calculation rests on an assumption about the number of independent regions that are potential targets of SDS across the genome. We assume there are 1,703 independent targets, following an estimate of the number of approximately independent LD blocks by Berisa and Pickrell (52). This is a conservative estimate of the target size because, for example, it only considers common genetic variation, such that numerous pairs of independent rare variants are likely included within the estimated haplotype blocks. The conservative estimate for the target size likely leads to an underestimate of the load. It is also important to note that using other values within the plausible range for our estimates of the mean selection coefficient and frequency of selection can result in very different load estimates. For example, with a selection coefficient of 0.005 and minor allele frequency of 1%, the mortality load increases to 1.7%. Nevertheless, our conclusion that the pervasive SDS we estimate in the genome need not generate a heavy mortality cost stands. Relatedly, a question that remains is when in the life cycle this mortality occurs. One possibility is *in utero* (57): embryonic survival rates in humans are reported to be as low as 50% (58) and neonatal sex differences in allele frequencies have been observed (59).

The Introduction noted the apparent disconnect between the abundant (and noncontroversial) evidence for sexually antagonistic selection acting on phenotypes vs. the limited (and controversial) evidence for SAS acting on the genome. This gap in our understanding is perhaps unsurprising for two reasons. First, the “selection in real time” strategy used here and elsewhere works with signals of selection resulting from just a single generation of selection, and so they are inherently very small. Second, many of the phenotypes in question are highly polygenic, so selection acting on individual sites underlying those traits will be very weak.

Our use of phased SNPs yields four benefits that advance the study sex-differential selection. First, phased SNPs increase power and decrease error relative to methods based on single SNPs (Figure 1C). We expect that additional power may be gained in future work by using haplotypes that include more than just two phased SNPs. (A technical challenge is that each additional SNP doubles the computational burden.) Second, we can detect sex-differential selection acting on SNPs that have not been genotyped. Third, allowing for selection on these “hidden” SNPs mitigates the underestimation of selection that results when one falsely assumes selection is acting on an observed SNP that is linked to the actual target (Figure 1C). Fourth, the number of genomic windows that we evaluate for sex-differential selection is several orders of magnitude smaller than the number of possible targets of SDS, so the statistical burden of multiple comparisons is dramatically mitigated.

What maintains the genetic polymorphisms that experience sex-differential selection? One contributing factor may be a life history tradeoff between viability and fecundity (Figure 3). Life history tradeoffs can help to maintain genetic variation for fitness (60), much as can sexually antagonistic fitness tradeoffs (2, 61). Sex-differential selection resulting from life history tradeoffs has been linked to loci of large effect (22, 24, 62) and to polygenic traits (3, 63, 64). Life history tradeoffs are not the only form of selection that can maintain polymorphisms subject to SDS. Theory shows that the range of parameters that maintains polymorphism is greatly expanded when alleles with sexually antagonistic effects show reversed dominance in females and males (2), and indeed dominance reversal has been observed (65, 66). Regardless of the details about how it does so, results from experimental evolution suggest SDS contributes to genetic variation for fitness-related traits (3, 67, 68). It is, however, crucial to keep in mind that some (perhaps most) genetic variation experiencing SDS may not involve stable polymorphisms. Indeed, Ruzicka *et al*. (40) found no evidence that SNPs under SDS in the human genome are subject to any form of balancing selection. Those polymorphisms might result from a variety of other evolutionary forces such as mutation and migration, or could in fact be transient (69).

A related puzzle is why SDS persists. In principle, each of the sexes could evolve to its optimum if appropriate genetic variation was available, for example in the form of variation in sex-specific (or sex differential) gene expression (70, 71). Two general kinds of hypotheses can be proposed. Selection may fluctuate in strength and direction rapidly enough that the sex-specific optima are never reached (68, 72). Alternatively, genetic constraints can cause allele frequencies and trait means in both sexes to evolve to suboptimal equilibria (73). Perfect genetic correlations (either positive or negative) between the expression of a trait in the two sexes would support the constraint hypothesis. But in fact this criterion is too stringent: some alleles that reduce the intersex correlation may be unconditionally deleterious, and the evolution of dimorphism in a given focal trait can be constrained by pleiotropy with other traits (15).

Several lines of evidence further support the constraint hypothesis. Artificial selection experiments on a flowering plant (74) and a fly (13) suggest there may be very limited genetic variation that would allow the evolution of increased sexual dimorphism. Estimates of the intersex genetic correlation for a variety of traits in several species (12, 75, 76) are very often near unity. In humans, large classes of allelic effects in one sex are a fixed multiple of their effects in the other (15), which implies there are strong constraints on the evolution of sex differences. The constraint hypothesis might be evaluated with phenotypic measures of selection on a trait such as body size in related species, which could determine if SDS acts in a consistent direction over appreciable evolutionary timescales. Given the limited evidence available, it seems plausible that SDS is an inescapable consequence of reproduction involving separate sexes.

## 4 Methods

### 4.1 UK Biobank samples and SNP quality controls

We used data from the UK Biobank (UKB), a large database with genetic and phenotypic information from over 500,000 participants in the UK. Individuals were categorized as female or male based on their sex chromosome karyotypes (XX vs XY). We removed outliers for missingness and heterozygosity, individuals with sex chromosome aneuploidy, and those individuals with a discrepancy between selfreported sex and genotypically inferred sex. Participants with high relatedness up to the 3rd degree were also excluded. To mitigate issues related to population structure, we kept only individuals identified as of ”white, British” ancestry based either on self-reported or genetic ethnicity data based on PCA. Lifetime reproductive success (LRS) was estimated using the reported number of live births for females and the recorded number of children fathered for males. We believe the number of children is a reasonable proxy as UKB participants range in age from 40-69. UK census data from 2006-2010, which represent the years of recruitment into the UKB, indicate that over 99% of women and 95% of men have their last child before age 45. Thus, we excluded individuals younger than 45 years of age. Moreover, there is a very strong genetic correlation between the number of offspring and number of grand-offspring in contemporary humans (77). The final dataset consisted of 303,824 individuals, including 163,959 females and 139,865 males.

We used only the 805,426 SNPs in the UKB that are genotyped and phased in the main analyses. We performed site-level QC procedures by removing SNPs that were not bi-allelic, had a minor allele frequency less than 1%, missing rates exceeding 5%, or exhibited excessive deviations from Hardy-Weinberg equilibrium. Several steps were taken to control for technical artifacts that could confound signals of SDS (See SI section A.2). After applying these filters, 554,944 SNPs remained. We also obtained a distribution of allele frequencies from UKB’s imputed data after applying the same filtering steps described above (on 9.9 million SNPs).

### 4.2 Novel method for detecting sex-differential selection

We used likelihood based on a population genetic model to estimate the selection coefficients pertaining to SDS. At autosomal sites, allele frequencies are expected to be equal in females and males at conception. SDS acting on a site causes allele frequencies in the sexes to diverge at that site. The frequencies also diverge at other sites in linkage disequilibrium with the target of SDS by hitchhiking (48). Our model seeks to detect this characteristic pattern. Here we outline the approach; full details are given in the SI. The likelihood of sampling the haplotypes observed in our sample is given by the multinomial distribution:

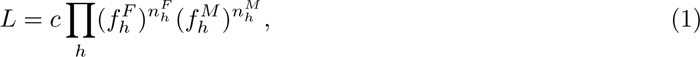

where 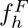 is the expected frequency of haplotype *h* in the population in females after selection, 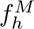 is the number of copies of that haplotype in our sample of females, 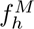 and 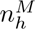 are the corresponding quantities for males, and *c* is a constant that is independent of SDS. The product ranges over all the haplotypes *h*. In our implementation, these consist of three sites: two observed SNPs and an unseen putative target of SDS that they flank. Assuming that these three sites are biallelic, there are 8 haplotype frequencies in each sex. Information about linkage disequilibria between the SNPs is captured by the haplotype frequencies.

We next express 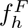 and 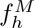 in terms of the strength of sex-differential viability selection and the haplotype frequencies at conception. We make the strong assumption that the selection coefficients for viability are sex-symmetric such that *s*_Female_ = *−s*_Male_ *≡ s_v_*. (SI section A.3 shows that when selection is not sex-symmetric, estimates of the selection coefficients can be biased either upwards or downward, but the relative magnitude of the bias is very small.) We assume Hardy-Weinberg equilibrium at conception; violations of this assumption have little effect on estimation unless they are extreme. Finally, we assume heterozygotes have intermediate fitness. If dominance is present, our estimates of *s_v_* represent the average effects of alleles on fitness rather than selection coefficients.

Under those assumptions, basic one-locus theory (60) shows that

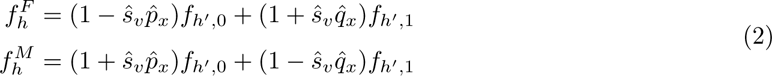

where *p_x_* is the frequency at conception of the minor allele at the unseen target of selection, *q_x_* = 1 *− p_x_*, and hats denote estimates. The quantity *f_h_′,*_0_ is the frequency at conception of the haplotype that carries allele 0 at the target and the set of alleles *h^′^*at the two flanking SNPs, while *f_h_′,*_1_ is the corresponding frequency with allele 1 at the target. The two terms on the right sides of Eqs. (2) average over the probabilities that the target carries allele 0 or allele 1. These expressions assume the population is sufficiently large that drift can be neglected.

To obtain expressions for *f_h_′,*_0_ and *f_h_′,*_1_, we need two quantities pertaining to the unseen target: the allele frequency *p_x_*, and the linkage disequilibria between that site and the observed SNPs that flank it. For *p_x_*, we chose the median minor allele frequency across the filtered subset of imputed SNPs (= 0.13), and showed with simulations that this value yields conservative (downward-biased) estimates of *s_v_*. Assuming larger values of *p_x_* tends to further underestimate *s_v_*, whereas assuming smaller values leads to overestimates (SI section B.1). Regarding linkage, we assume the target is midway between the flanking SNPs; simulations show the results are surprisingly robust to violations of this assumption. The SI (section A.3) gives further details.

We obtained estimates for the SDS viability selection coefficient for each pair of adjacent phased SNPs in the dataset. The maximum likelihood estimate *ŝ_v_* was obtained by substituting Eqs. (2) into Eq. (1), then maximizing *L* numerically with respect to *ŝ_v_*.

Selection coefficients for total lifetime reproductive success (comprising both viability and fertility selection) were estimated by weighting the haplotype frequencies and allele counts by the average fecundities of individuals carrying those haplotypes and alleles. These estimates, denoted *ŝ_T_*, were in turn used to estimate selection coefficients for fecundity selection with the relation Details are given in SI section A.3.

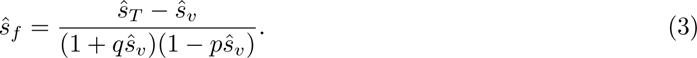

We used simulation to assess the performance of our method. We generated haplotypes by sampling the minor allele frequencies and linkage disequilibria between pairs of adjacent phased SNPs in the UKB. Next we generated an unseen target of SDS between these flanking SNPs. In baseline simulations, we assumed that the minor allele frequency at this target was 0.13, and that the target was midway between the flanking SNPs. We simulated selection of varying strengths acting on this target, generated samples of haplotypes in females and males after selection acts, then ran these pseudodata through our estimation pipeline. To evaluate the robustness of the model to violations of its assumptions, we ran simulations with different minor allele frequencies at the target, and allowed it to lie at different positions between the observed flanking SNPs.

### 4.3 Estimating SDS across the genome

Analyses were performed in two-site sliding windows across the genome. For each window, we estimated sex-differential viability selection coefficients using likelihood (Eq. 1), and calculated per-site standard errors and *p* values. To estimate the strength of SDS acting across the lifetime and on fecundity, we calculated projected haplotype counts in offspring using the recorded number of children for each participant given by the UKB. To generate null distributions, we permuted the sex labels in the dataset once and refit the likelihood models. We pruned our results for LD when performing all downstream analyses (See SI section A.4).

### 4.4 Linking SDS to sex-specific allelic effects

To investigate the impact of SDS on complex traits, we developed a model that links the strength of selection (*s*) and the additive effects of a biallelic locus (*β_f_* and *β_m_*) in females and males on phenotypes. We followed a model developed in Zhu *et al*. (15) that assumes equal allele frequencies at conception under symmetric sexually-antagonistic selection, and assumes a linear relation between *s* and *β*. These assumptions lead to the relationship:

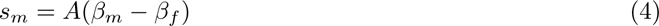

(see equation (SI 20)). Here *A* represents the intensity of sexually antagonistic selection on the focal trait (see equation (SI 21)).

We used sex-specific SNP effect sizes from Zhu *et al*. (15) on 27 quantitative traits with estimates of SNP heritability greater than 7.5%. Their estimates were obtained through sex-stratified GWAS and adjusted using multivariate adaptive shrinkage (*mash*). To estimate *A*, for each of the 27 traits we performed a weighted standard major axis regression for all three modes of selection. This approach accounts for uncertainty in both the selection gradients and trait effect sizes, using the standard errors for the selection coefficients and the standard deviations provided by *mash*. To mitigate the influence of linkage disequilibrium (LD) between sites, we divided the genome into the 1,703 approximately independent haplotype blocks identified by Berisa and Pickrell (52). We performed the regressions by sampling one SNP from each haplotype block, resulting in 1,689 SNPs with their corresponding estimated selection coefficients and sex-stratified marginal effect sizes for each trait. This procedure was repeated 1,000 times per trait. The mean slope of the 1,000 regressions was divided by the standard deviation of the slopes to obtain *Z* -scores for sexually antagonistic selection.

### 4.5 Estimating the intensity and frequency of SDS

We used Approximate Bayesian Computation (ABC) to estimate the average selection coefficient, *ŝ*, and the fraction of haplotype windows under selection, *F* . Values of each parameter were drawn from a prior distribution. We then simulated the resulting haplotype frequencies, and ran these pseudodata through our estimation pipeline to obtain estimates of *ŝ* and *F* . The procedure was repeated 50,000 times, and the 1% of estimates of *ŝ* and *F* that best matched the true values were retained as posterior estimates (SI figure B.8.1). Outcomes were robust to the rejection threshold and number of simulations (SI figures B.8.2 and B.8.3). The mortality load was then calculated as a function of allele frequency using these posterior estimates. Details are given in SI section A.8.

## 5 Acknowledgements

We thank Pavitra Muralidhar, Filip Ruzicka, and the two reviewers for providing helpful comments on the manuscript. This work was supported by NIH grant GM11685307 to M.K., and NIH grant GM151108 and a Pew Scholarship to A.H. This project was conducted using the UK Biobank resource under application number 61666.

## 6 Data availability

All of the data generated by this study can be found at Zenodo (https://doi.org/10.5281/zenodo.11992199) and all original code is publicly available at https://github.com/harpak-lab/SDS humans.

## Notes

https://doi.org/10.5281/zenodo.11992199

https://github.com/harpak-lab/SDS_humans

### Competing Interest Statement

The authors have declared no competing interest.

