## Supplemental Information (SI) Appendix for "The battle of the sexes in humans is highly polygenic"

### Section A: Supplemental Methods

#### A.1. UK Biobank samples

The UK Biobank (UKB) is a large human database containing detailed genetic, phenotypic and offspring data for over 500,000 participants of both sexes between 40 and 69 years of age in the United Kingdom (1). We leveraged participant metadata to perform several individual-level quality controls. Namely, we removed individuals that were identified as outliers for missingness and heterozygosity (UKB field 22027), that exhibited sex chromosome aneuploidy (field 22019), or that showed a mismatch between self-reported sex and genotypically inferred sex (fields 31 and 22001 respectively). Further, we excluded individuals with relatedness up to the 3rd degree (field 22020). To control for effects of population structure, we removed samples not identified of "white British" ancestry in either the self-reported or the genetic ethnicity fields (field 22006).

We extracted "number of live births" for females (field 2734) and "number of children fathered" for males (field 2405) and took the maximum number of offspring reported across multiple assessment periods as a proxy for lifetime reproductive success (LRS) for a given participant. Following Ruzicka *et al.* (2), we excluded individuals with unreported LRS, who were under 45 years of age at recruitment (field 21022), who reported a larger number of offspring at earlier assessments than later ones, and who reported 20 or more offspring. Participants who had withdrawn from the database by the time of this study were also excluded.

#### A.2. UK Biobank genotype data and site-level quality controls

The UKB contains 805,426 array-genotyped, quality-controlled SNPs (1). Two genotype arrays, which share over 95% of common variants, were used to call SNPs. Approximately 450,000 individuals were genotyped on the custom UK Biobank Affymetrix Axiom array and approximately 50,000 individuals genotyped using the UK BiLEVE array. We focused only on these array-genotyped SNPs since they have been statistically phased onto chromosomes, which is crucial for our analyses. We performed additional site-level quality controls using both PLINK v1.9 and PLINK2 (3): we removed sites that had more than two alleles, minor allele frequencies less than 1%, missing rates exceeding 5%, and excessive deviations from Hardy-Weinberg ($p < 10^{-6}$, exact test; (2, 4)).

In addition to basic site-level quality control measures, we took several steps to minimize bioinformatic and technical artifacts that can generate spurious signals of sex-differential selection. High sequence similarity between the autosomes and sex chromosomes can result in the mismapping of Y-linked reads to autosomal regions (5, 6) and mis-hybridization of autosomal sequences to sex chromosomes (7, 8). To minimize these artifacts, we first filtered out sex-differential variants from the data by removing SNPs that were monoallelic in either males or females. Second, we removed sites with a deficit of minor allele homozygotes, significant differences in missingness between males and females, or excessive heterozygosity (2, 8). Third, we removed all SNPs that exhibited known homology with a sex chromosome region identified in the same dataset by Kasimatis *et al.* (8) as well as those identified in Galichon *et al.* (9). After estimation, we filtered our dataset further by excluding adjacent pairs of sites where the $r^2$ between them was less than 0.1 and where either sex had haplotype counts less than 10. We also filtered out sites located in the major histocompatability complex (MHC; chr6: 28,477,797 - 33,448,354) to avoid issues due to mismapping. This left us with 248,059 windows.

#### A.3. The likelihood method

##### Estimating the targets and strength of SDS using likelihood

Assuming that individuals are sampled randomly from the population, the log likelihood of obtaining the observed frequencies of haplotypes is determined by the multinomial distribution:

$$ln(L) = c + \sum_h \Big\{ n_h^F ln(f_h^F) + n_h^M ln(f_h^M) \Big\}, \tag{SI 1}$$

where $f_h^F$ is the expected frequency of haplotype $h$ in the population in females after selection, $n_h^F$ is the number of copies of that haplotype in our sample, $f_h^M$ and $n_h^M$ are the corresponding quantities for males, and $c$ is a constant that is independent of SDS and so does not affect estimation. The sum ranges over all the haplotypes $h$. In our implementation, these consist of two observed SNPs flanking an unseen putative target of SDS. Assuming that these three sites are biallelic, there are 8 haplotype frequencies in each sex. Information about linkage disequilibria between the SNPs is captured by the haplotype frequencies.

We next express $f_h^F$ and $f_h^M$ in terms of the strength of sex-differential viability selection and the haplotype frequencies at conception. We assume that the selection coefficients for viability are sex-symmetric such that $s_{Female} = -s_{Male} \equiv s_v$, and that zygotes are in Hardy-Weinberg equilibrium (HWE) at conception. Unless they are extreme, violations of these assumptions have little effect on estimates of $s_v$ (see below). Finally, we assume heterozygotes have intermediate fitness. If dominance is present, $s_v$ represents the additive effect of the allele on fitness rather than the selection coefficient.

Under those assumptions, basic one-locus theory (10) shows that

$$\begin{aligned} f_h^F &= (1 - \hat{s}_v \hat{p}_x) f_{h',0} + (1 + \hat{s}_v \hat{q}_x) f_{h',1}, \\ f_h^M &= (1 + \hat{s}_v \hat{p}_x) f_{h',0} + (1 - \hat{s}_v \hat{q}_x) f_{h',1}, \end{aligned} \tag{SI 2}$$

where $p_x$ is the frequency at conception of the minor allele at the unseen target of selection $x$, $q_x = 1 - p_x$, and hats denote estimates. The quantity $f_{h',0}$ is the frequency at conception of the haplotype that carries allele 0 at the target and the set of alleles $h'$ at the two flanking SNPs; $f_{h',1}$ is the corresponding frequency with allele 1 at the target. The two terms on the right sides of Eqs. (SI 2) average over the probabilities that the target carries allele 0 or allele 1, and they neglect terms that are $O(\hat{s}_v^2)$.

Next, we write the haplotype frequencies in terms of allele frequencies and linkage disequilibria. We denote the flanking SNP to the left of the unseen target as site 1 and that to the right at site 0. Haplotypes that carry alleles (0, 0), (0, 1), (1, 0), and (1, 1) at the flanking SNPs are denoted respectively

$h' = 0, 1, 2, 3$. Then

$$
\begin{aligned}
f_{0,0} &= q_0 q_1 \hat{q}_x + \hat{q}_x D_{1,0} + q_0 D_{1,x} + q_1 D_{x,0},\\
f_{1,0} &= p_0 q_1 \hat{q}_x - \hat{q}_x D_{1,0} + p_0 D_{1,x} - q_1 D_{x,0},\\
f_{2,0} &= q_0 p_1 \hat{q}_x - \hat{q}_x D_{1,0} - q_0 D_{1,x} + p_1 D_{x,0},\\
f_{3,0} &= p_0 p_1 \hat{q}_x + \hat{q}_x D_{1,0} - p_0 D_{1,x} - p_1 D_{x,0},\\
\\
f_{0,1} &= q_0 q_1 \hat{p}_x + \hat{p}_x D_{1,0} - q_0 D_{1,x} - q_1 D_{x,0},\\
f_{1,1} &= p_0 q_1 \hat{p}_x - \hat{p}_x D_{1,0} - p_0 D_{1,x} + q_1 D_{x,0},\\
f_{2,1} &= q_0 p_1 \hat{p}_x - \hat{p}_x D_{1,0} + q_0 D_{1,x} - p_1 D_{x,0},\\
f_{3,1} &= p_0 p_1 \hat{p}_x + \hat{p}_x D_{1,0} + p_0 D_{1,x} + p_1 D_{x,0},
\end{aligned}
\tag{SI 3}
$$

where $D_{i,j}$ is the linkage disequilibrium between sites $i$ and $j$. These expressions assume there is no three-way disequilibrium between the sites.

Eqs. (SI 3) involve two unobserved nuisance variables pertaining to the unseen target of selection: the allele frequency $p_x$, and the linkage disequilibria between that site and the observed flanking SNPs. We set $p_x = 0.13$, a value that is similar to the median minor allele frequency across all the SNPs in our filtered dataset of the imputed genotypes from Bycroft *et al.* (1). Simulations presented in the next section show it yields conservative (downward-biased) estimates of $s_v$. Regarding linkage, we assume the target is midway between the flanking SNPs in the sense that

$$r_{1,0} = r_{1,x} r_{x,1}, \tag{SI 4}$$

where $r_{i,j}$ is the correlation between allelic states (0 or 1) at sites $i$ and $j$. That relation implies

$$
\begin{aligned}
D_{1,x} &= \Big(|D_{1,0}|\hat{p}_x \hat{q}_x\Big)^{1/2} \Big(\frac{p_1 q_1}{p_0 q_0}\Big)^{1/4},\\
D_{x,0} &= I(D_{1,0})\Big(|D_{1,0}|\hat{p}_x \hat{q}_x\Big)^{1/2} \Big(\frac{p_0 q_0}{p_1 q_1}\Big)^{1/4},
\end{aligned}
\tag{SI 5}
$$

where $|.|$ denotes the absolute value and $I(D_{1,0}) = -1$ if $D_{1,0} < 0$, or $+1$ if not.

Substituting Eqs. (SI 2 – SI 5) into (SI 1) gives an expression for the log likelihood as a function of the estimated selection coefficient for viability, $\hat{s}_v$. The maximum likelihood estimate is obtained by maximizing it numerically with respect to $\hat{s}_v$.

#### Violations of model assumptions

We used both analytic methods and simulations to determine the sensitivity of our estimation framework to violations of the assumptions. First consider the assumption that viability selection is symmetric in the sexes such that the selection coefficients are $s_{Female} = -s_{Male} \equiv s_v$. Consider a site that is the target of selection. If SDS truly is sex-symmetric, one can show that the maximum likelihood estimator for the selection coefficient is

$$\hat{s}_v = \frac{2N^F N^M (n_1^F n_0^M - n_0^F n_1^M)}{2n_0^F n_1^F (N^M)^2 + 2n_0^M n_1^M (N^F)^2 + (n_0^F n_1^M - n_1^F n_0^M)^2}, \tag{SI 6}$$

where $n_i^F$ is the number of copies of allele $i$ in females, and $N^F = n_0^F + n_1^F$ is the total number of copies in females; $n_i^M$ and $N^M$ are the corresponding quantities in males. Now imagine for simplicity that there is no sampling variation and so the number of copies of alleles in the sample is proportional to their frequencies after selection. Then with symmetric SDS,

$$\begin{aligned} n_0^F &= (1 - s_v p_x) q_x N^F, \\ n_1^F &= (1 + s_v q_x) p_x N^F, \\ n_0^M &= (1 - s_v p_x) q_x N^M, \\ n_1^M &= (1 + s_v q_x) p_x N^M. \end{aligned} \tag{SI 7}$$

Substituting these quantities into (SI 6) yields $\hat{s}_v = s_v$, which shows the estimator is unbiased. Now consider an alternative situation in which selection is restricted entirely to one sex, say females, such that $s_{Female} = 2s_v$ and $s_{Male} = 0$. (The factor of 2 appears in the expression for $s_{Female}$ so that the difference between the selection coefficients in the two sexes is equal to the case of symmetric selection.) Then the expected numbers of allele copies in the sample are

$$\begin{aligned} n_0^F &= (1 - 2s_v p_x) q_x N^F, \\ n_1^F &= (1 + 2s_v q_x) p_x N^F, \\ n_0^M &= q_x N^M, \\ n_1^M &= p_x N^M. \end{aligned} \tag{SI 8}$$

Substituting these quantities into (SI 6), we find that the relative bias in the estimator is

$$bias = (1 - 2p_x) s_v \tag{SI 9}$$

(to leading order in $s_v$). Thus assuming that SDS is sex-symmetric when in fact it is sex limited results in a biased estimate of the selection coefficient. This results shows that the relative bias, which can be positive or negative, can be no larger than $s_v$. As we expect selection coefficients to be typically much less than unity, the relative bias induced by violating the sex symmetry assumption is expected to be small.

##### Performance of the haplotype method under simulation

We used simulations to evaluate our assumptions regarding the minor allele frequency and linkage disequilibria involving the unseen target of selection. Haplotype frequencies were obtained following equation (SI 3). Minor allele frequencies for the two flanking sites ($p_1$ and $p_0$) and their corresponding values for linkage disequilibrium (in terms of the correlation between them, $r_{1,0}$) were sampled from the filtered distribution of minor allele frequencies in the phased, genotyped dataset. The minor allele frequency for the unknown target ($p_x$) was sampled from the filtered distribution of minor allele frequencies in the imputed dataset (> 9 million SNPs, using the criteria of section A.2). We varied the location of the target site within each haplotype by setting the distance of the of the target from site 0 to

$$d = \frac{ln(r_{x,0})}{ln(r_{1,0})}, \tag{SI 10}$$

where $d = 0$ if the target is completely linked to site 0, $d = 1$ if it is completely linked to site 1, and $d = 0.5$ if it lies midway between the flanking sites. The disequilibria between the target and the flanking sites, $D_{1,x}$ and $D_{x,0}$, are found by substituting (SI 10) into (SI 5). In turn, those quantities are substituted into (SI 3) to obtain the haplotype frequencies in zygotes. The expected frequencies in adults following selection are then given by Equations (SI 2). We simulated datasets by sampling the multinomial distribution using those frequencies, with a sample size of $N = 300,000$ haplotypes for each sex. For each dataset, an estimate of the selection coefficient was obtained using the likelihood method described by Equations (SI 1 - SI 6), assuming that selection is sex-symmetric, that the minor allele frequency at the target is $p_x = 0.13$, and that the target is midway between the flanking sites ($d = 0.5$). We varied $d$ from 0 to 0.5 and the selection coefficient between $s_v = 10^{-3}$ and $10^{-1}$. Each parameter combination was replicated $10^5$ times.

The results are shown in Figure 1C. We see that the estimate of $s_v$ is very little affected by the position of the target.

Next consider our assumption that the minor allele frequency at the unseen target is $p_x = 0.13$. (Recall that value was chosen because it is close to the median minor allele frequency among the filtered, imputed SNPs in the UK Biobank dataset.) We again simulated datasets as described above, but now allowed the true value of $p_x$ to vary. The results show that assuming $p_x = 0.13$ yields the best performance (Figures B.1.1 - B.1.3). Consistent with intuition, larger values of $p_x$ lead to underestimates of $s_v$ (because the same change in an allele frequency results from weaker selection when the allele frequency is closer to $\frac{1}{2}$), while smaller values of $p_x$ have the converse effect.

##### The likelihood function for fecundity selection and total selection

We can also use the log likelihood function (SI 1) to estimate selection coefficients pertaining to lifetime fitness, that is, comprising the effects of SDS on both viability and fecundity. Again assuming that selection is sex-symmetric, we simply weight the haplotype counts in each sex by the average fecundity of individuals that carry those haplotypes and alleles. The weighted haplotype counts for females are

$$\tilde{n}_h^F = \frac{W_h^F n_h^F}{\bar{W}^F}, \tag{SI 11}$$

where

$$W_h^F = \frac{1}{2} c_h^F, \tag{SI 12}$$

where $c_h^F$ denotes the cumulative number of recorded offspring for females with haplotype $h$, and

$$\bar{W}^F = \frac{1}{N^F} \sum_h W_h^F \tag{SI 13}$$

where $N^F$ denotes the number of female haplotypes. Proceeding as described in the previous section then yields estimates of the total selection coefficients for SDS, denoted as $\hat{s}_T$. To estimate a selection coefficient for fecundity, written $\hat{s}_f$, we write the change in the allele frequency from the the start of the current generation to the start of the next as the sum of the changes caused by viability and fecundity selection:

$$\Delta_T p = s_v pq + s_f p'q', \tag{SI 14}$$

where primes denote frequencies after viability but before fecundity selection (in the adults). Then

$$\Delta_T p = s_v pq + s_f p(1 + s_v q)q(1 - s_v p), \tag{SI 15}$$

which on rearranging gives

$$s_f = \frac{s_T - s_v}{(1 + qs_v)(1 - ps_v)}. \tag{SI 16}$$

Substituting the estimated selection coefficients $\hat{s}_v$ and $\hat{s}_T$ for the corresponding population parameters gives Eq. 3 of the main text.

#### Identifying targets of SDS

To evaluate which regions in the genome may be targets of SDS, we calculated standard errors for each per-window estimate of the selection coefficient using a parametric bootstrap. For each window, samples of haplotypes were simulated by sampling from the observed haplotypes with replacement, from which we estimated the selection coefficient as described above. For each window in the dataset, we repeated this procedure 100 times, calculated the standard error of $\hat{s}$, and used the resulting $Z$-scores to calculate the approximate significance ($p$ value). We validated these results by comparing them against the likelihoods, and found the two methods are broadly consistent.

#### Comparison with Ruzicka et al.

Ruzicka *et al.* (2) recently analyzed the UK Biobank dataset using different statistical methods. We wished to determine if our approach gives comparable results. Following earlier studies (eg, (11)), Ruzicka et al. sought to detect SDS using $F_{\mathrm{ST}}$ to quantify allele frequency differences between the sexes. That statistic can be related to the selection coefficients for SDS via Eq. (5) in Cheng and Kirkpatrick (11):

$$s = \sqrt{\frac{F_{\mathrm{ST}}}{pq}}. \tag{SI 17}$$

To compare the results in their study and ours, we estimated selection coefficients on all the genotyped SNPs (LD-pruned and filtered as discussed in A.2 and A.4 below) using the estimators given in equation (SI 6) for total and viability selection and equation (SI 16) for fecundity selection. A total of 190,374 SNPs overlapped our datasets. We converted their $F_{\mathrm{ST}}$ metrics for viability, fecundity and total selection (respectively designated as adult, reproductive and gametic $F_{\mathrm{ST}}$ in their study) into selection coefficients using equation (SI 17), then calculated Pearson's correlation coefficients between those values and our estimates (SI figure B.5.1).

#### A.4. Quantifying signals of sex-differential selection in the UK Biobank dataset

To quantify signals of SDS, we fit the likelihood models on the filtered dataset of 554,944 phased SNPs (see filtering steps above). We fit models in sliding windows of two SNPs at a time (see Figure 1), a step size of one SNP. With a haplotype length of two and only considering biallelic sites, there are 4 possible observable haplotypes in each window. In total, we analyzed 550,725 windows.

To estimate sex-differential viability selection coefficients, we fit equation (SI 1) to each window as described above. To calculate standard errors of the estimates for each site, we employed a parametric bootstrap procedure using multinomial sampling. This approach allowed us to generate a distribution of bootstrapped selection coefficients by sampling haplotype counts under a multinomial model given the haplotype frequencies in the observed data and fitting the likelihood model. For each window, we performed this bootstrapping procedure 100 times. Standard errors were then estimated by calculating the standard deviations of the 100 bootstrapped estimates for each site. Per-site $Z$-scores were also calculated by dividing each site's estimated selection gradient by its standard error, and the $Z$-scores used to calculate per-site $p$ values based on the standard normal distribution.

For estimation of total SDS (the combination of viability and fecundity selection), we calculated the LRS-weighted haplotype counts before fitting the model (see equation SI 11). To calculate standard errors, we performed a modified parametric bootstrap of the procedure described above for viability selection. We modified the procedure by first constructing a matrix of haplotype counts for each window for both sexes, with 4 rows representing all possible two-site haplotypes and each column representing the recorded LRS values. We generated a bootstrapped haplotype matrix under a multinomial model given the observed haplotype frequencies within each cell of the matrix, and fit the likelihood model. We perform this procedure 100 times per window in each sex to obtain bootstrapped distributions of the estimated selection coefficients. Standard errors, $Z$-scores, and $p$ values are calculated the same way as for the viability selection coefficients.

To estimate sex-differential fecundity selection, we calculated selection coefficients according to equation (SI 16). To estimate standard errors, we perform similar parametric bootstrapping as described for total selection, employing the same bootstrapped matrix to obtain both the bootstrapped adult haplotype counts as well the bootstrapped projected haplotype counts in the offspring. These counts can be used to get bootstrapped values for both viability and total selection coefficients that allow us to obtain bootstrapped estimates of the fecundity selection coefficients. Standard errors, $Z$-scores, and $p$ values are estimated using the same calculation as for viability and fecundity estimates.

We generated null distributions of the selection coefficients for all three modes of selection by permuting the sex labels once and refitting the likelihood models. Standard errors, $Z$-scores, and $p$ values were calculated on the permuted datasets in the same way as in the observed datasets, following the procedures outlined above. We pruned the dataset for LD using the first site in each window using PLINK (3) at a threshold of $r^2 >= 0.2$ (flag –indep-pairwise 50 10 0.2) before comparing our null distributions to our observed distributions. After filtering, there were 127,834 windows left for that analysis.

To test for an excess of large selection coefficients, we first compared our observed estimates for each mode of selection with their respective empirical nulls using Mann-Whitney $U$ tests. Second, we binned the viability, fecundity, and total estimates by absolute value and performed $\chi^2$ tests on the difference in the number of values in each bin between the observed and null distributions. The results are shown in Figure 2.

#### A.5. Estimating the proportion of sites under sexually antagonistic fecundity selection

To separate concordant from antagonistic SDS acting on fecundity, we modified equation (SI 6) to estimate fecundity selection on each sex separately by using allele counts before fecundity selection (in adults) and allele counts post selection (allele counts adjusted for LRS), as described in section A.3. This calculation was done on both the observed and permuted haplotype counts. In females, the fecundity selection coefficient becomes

$$s_f^F = \frac{\tilde{n}_1^F(1-p^F) - \tilde{n}_0^F p^F}{(\tilde{n}_0^F + \tilde{n}_1^F)(1-p^F)p^F}, \qquad \text{(SI 18)}$$

where $\tilde{n}_i^F$ is the LRS-adjusted count for allele $i$ in females, and $p^F$ is the frequency of allele 1 in adult females. An analogous expression gives $s_f^M$ for males. Because the selection coefficients are signed and sex-specific, we find the whether a site is under antagonistic selection by calculating the product:

$$s_f^* = s_f^F \times s_f^M. \qquad \text{(SI 19)}$$

Negative values of $s_f^*$ indicate antagonistic selection. We took the absolute values for all the negative $s_f^*$ and binned them into 100 quantiles based on values calculated on the permuted data. We used $\chi^2$ tests to assess the relative excess at the top 1% of these values (see SI figure B.6.1).

#### A.6. Functional categorization of sex-differentiated alleles

To annotate potential functional consequences of variants under SDS, we employed Ensembl's Variant Effect Predictor (VEP) for build GRCh37 (12) on SNPs located in significant windows (after sequential Bonferroni correction, $\alpha = 0.05$) for viability and fecundity selection. We obtained genes with the closest protein-coding transcription start site within 5 kb upstream and downstream of each variant (using the –nearest flag, SI table B.4.1).

#### A.7. Relating sexually-antagonistic selection to complex traits

In order to investigate the effect of SDS on complex traits, we first formulated a model that relates the strength of selection ($s$) in a window and the additive effects of a biallelic locus($\beta_f$ and $\beta_m$) in females and males on phenotypes. As with the likelihood approach, we assume equilibrium in allele frequencies under symmetric sexually-antagonistic selection. We follow a model developed in Zhu *et al.* (13) that relates sex differences in trait effect size to selection coefficients ($s_f = a_f\beta_f$ in females, $s_m = a_m\beta_m$ in males), where $a_f$ and $a_m$ are the intensity of sexually antagonistic selection on alleles affecting a trait in females and males, respectively. We find that

$$s_m - s_f = 2A(\beta_m - \beta_f), \tag{SI 20}$$

where

$$A = \frac{a_m a_f}{a_m + a_f}. \tag{SI 21}$$

With symmetric sexually antagonistic selection, $s_f = -2s_m$, and so

$$s_m = A(\beta_m - \beta_f). \tag{SI 22}$$

This equation provides a linear relationship between the strength of selection in males and the sex difference in effects on phenotypes.

To obtain estimates of SNP effect sizes on traits, we used data from Zhu *et al.* (13) on 27 quantitative traits that were chosen based on their relatively high SNP heritabilities ($> 7.5\%$). These per-SNP effect size estimates were calculated via a sex-stratified GWAS and further adjusted using multivariate adaptive shrinkage (*mash*). Marginal SNP effect sizes were polarized so that both SNP effects on traits and selection coefficients act on the same allele (allele 1).

To estimate the intensity of sexually antagonistic selection on these traits ($A$), we conducted a weighted, standard major axis (SMA) regression (14, 15) for all three modes of selection. We used SMA regression to account for uncertainty in estimates of both variables (selection coefficients and trait effect sizes). We performed the regressions with a weighting, $v_i$, that is inversely proportional to the sum of the variances of our estimates of selection and the effect sizes at site $i$ as

$$v_i = \frac{1}{Var(\hat{s}_i) + Var(\beta_{m,i} - \beta_{f,i})}, \tag{SI 23}$$

where

$$Var(\hat{s}_i) = SE[\hat{s}_i]^2, \tag{SI 24}$$

and

$$Var(\beta_{m,i} - \beta_{f,i}) = SD[\beta_{f,i}]^2 + SD[\beta_{m,i}]^2, \tag{SI 25}$$

in which $SE[\hat{s}_i]$ is the standard error for the selection coefficient at site $i$, which we estimate via bootstrap (see SI section A.4 for details) and $SD[\beta_{m,i}]$ and $SD[\beta_{f,i}]$ are the estimated standard deviations for the *mash*-reduced trait effect sizes for males and females reported by Zhu et al (13) at site $i$. With $n$ sites, the individual weightings become

$$w_i = \frac{v_i}{\sum_i v_i}. \tag{SI 26}$$

To reduce the influence of LD between sites, we subdivided the genome into the 1,703 approximately independent haplotype blocks for Europeans given by Berisa and Pickrell (16). We performed the above regressions by sampling one window per block and taking the effect sizes for the SNP with the lowest local false sign rate (LFSR, reported in (13)). We thus obtained 1,689 SNPs per regression along with their associated estimated selection coefficients and the sex-specific marginal effect sizes. The procedure was iterated 1,000 times for each trait, and the mean slope value was divided by the standard deviation of the slopes to obtain $Z$-scores for sexually antagonistic selection. The resulting $Z$-scores were converted into $p$ values and were FDR corrected using the Benjamini-Hochberg procedure.

#### A.8. Estimating the mortality load resulting from sex-differential viability selection

We estimated the strength ($s$) and frequency ($F$) of SDS by simulating selection across 3,406 sites in the human genome and comparing them to our observed values. We randomly chose two of the adjacent phased SNPs from each haplotype block. For each of these, we sampled 50,000 pairs of values of $s$ and $F$. Prior values of $F$ were distributed such that $\log_{10}(F)$ was uniformly distributed on [-4, 0]. Given a value of $F$, the value of $s$ was 0 with probability $1 - F$ and otherwise was distributed such that $\log_{10}(s)$ was uniformly distributed on [-5, 1]. Given $s$, samples of female and male haplotypes after selection were simulated as described above (Performance of the haplotype method under simulation). These pseudodata were used to estimate $s$ with the likelihood model, giving a bivariate probability density function (PDF) in $s$ and $F$. The agreement between this PDF and that estimated from the real data was measured by the sum of squares. Approximate Bayesian Computation as implemented in the *abc* library in R (17) was used with the rejection method, retaining the best 1% (= 500) values of $s$ and $F$ as the posterior distribution (SI figure B.8.1). We found that the results were insensitive to both the number of posterior values retained and changes in the number of prior values simulated (SI figures B.8.2 and B.8.3). We took the maximum density of the joint posterior distribution to yield point estimates for $s$ and $F$ and calculated the 90% high-density credible intervals for these estimates using their posterior distributions.

Assuming multiplicative selection, we estimate the mortality load (18) as

$$L = 1 - (1 - sp)^{nF}, \qquad \text{(SI 27)}$$

where $s$ is the point estimate for the viability selection coefficient, $F$ is the point estimate for the frequency of SDS, $p$ is the allele frequency at the target of selection, and $n$ is the number of potential targets of SDS. We assume the number of potential targets are the 1,703 approximately independent LD blocks in Europeans (EUR) estimated by Berisa and Pickrell (16).

### Section B: Supplemental Figures and Tables

#### B.1. Simulation results

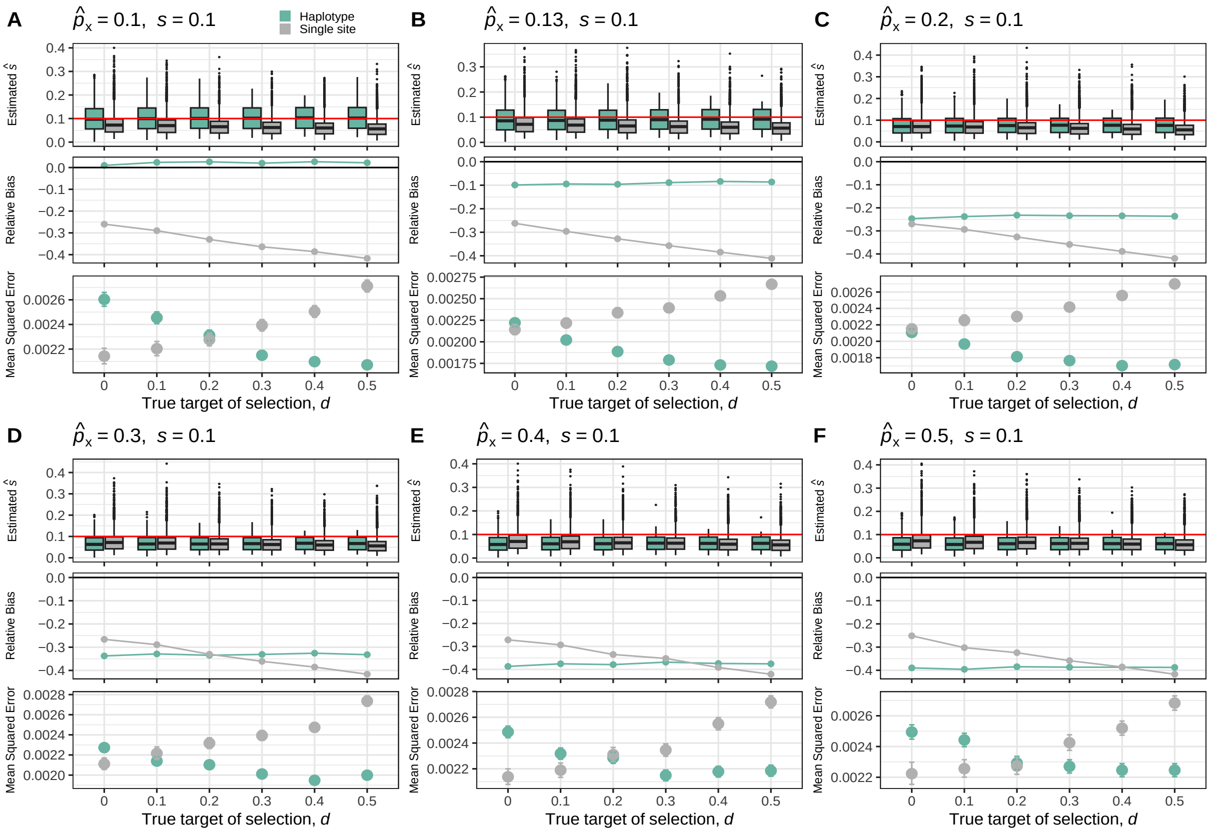


**Figure B.1.1:** Simulation results when $s = 0.1$. Panels A-F show the distribution of estimated selection coefficients (top pane), mean relative bias (middle pane), and mean squared errors (bottom pane) for different values of $\hat{p}_x$ . The red line in the top pane indicates the true value of $s$.

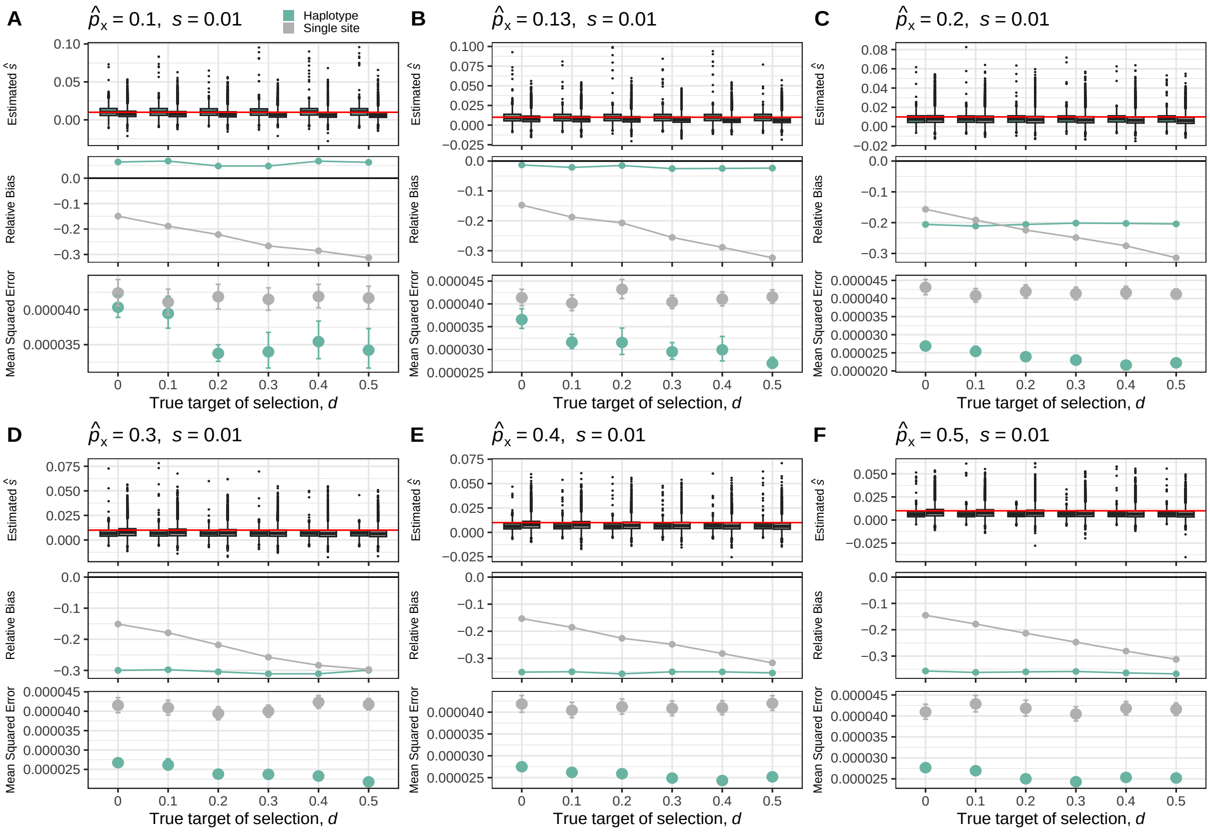


**Figure B.1.2:** Simulation results when $s = 0.01$. Panels A-F show the distribution of estimated selection coefficients (top pane), mean relative bias (middle pane), and mean squared errors (bottom pane) for different values of $\hat{p}_x$ . The red line in the top pane indicates the true value of $s$.

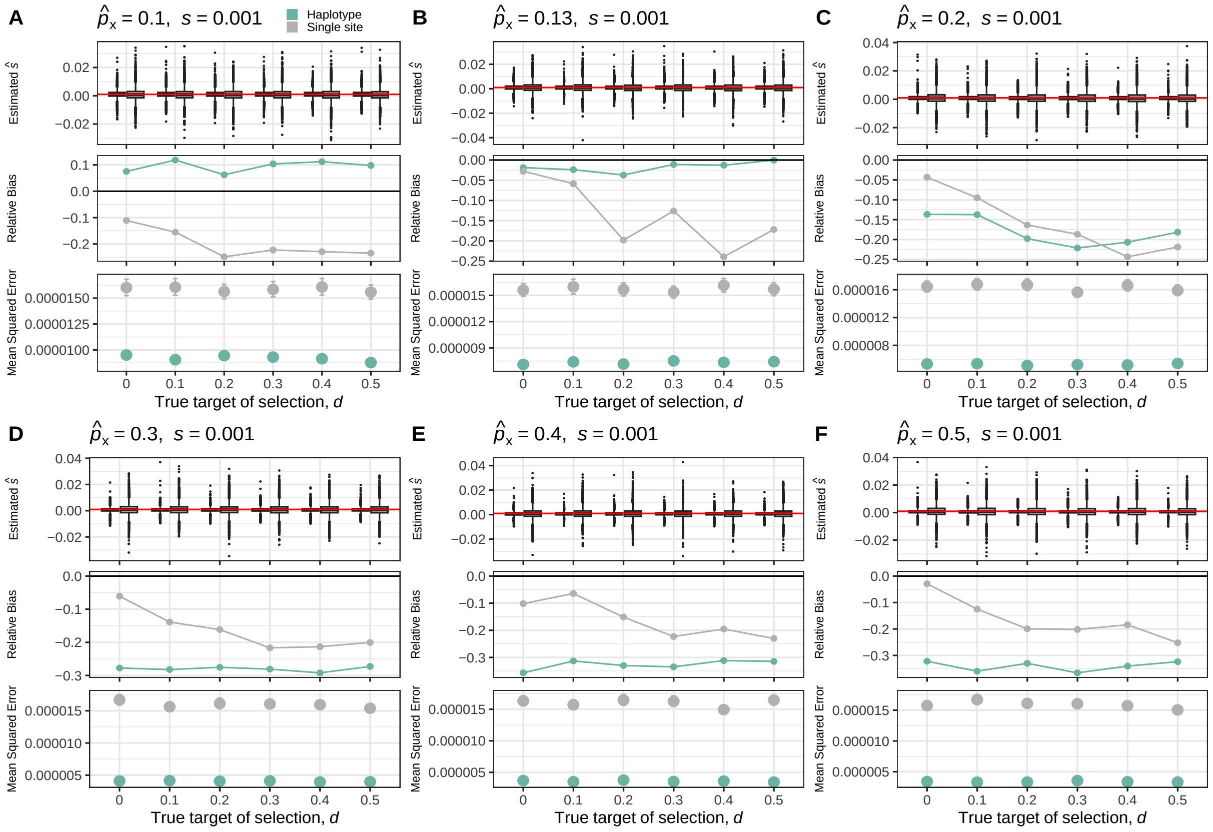


**Figure B.1.3:** Simulation results when $s = 0.001$. Panels A-F show the distribution of estimated selection coefficients (top pane), mean relative bias (middle pane), and mean squared errors (bottom pane) for different values of $\hat{p}_x$ . The red line in the top pane indicates the true value of $s$.

#### B.2. Empirical and permuted distributions of $\hat{s}$

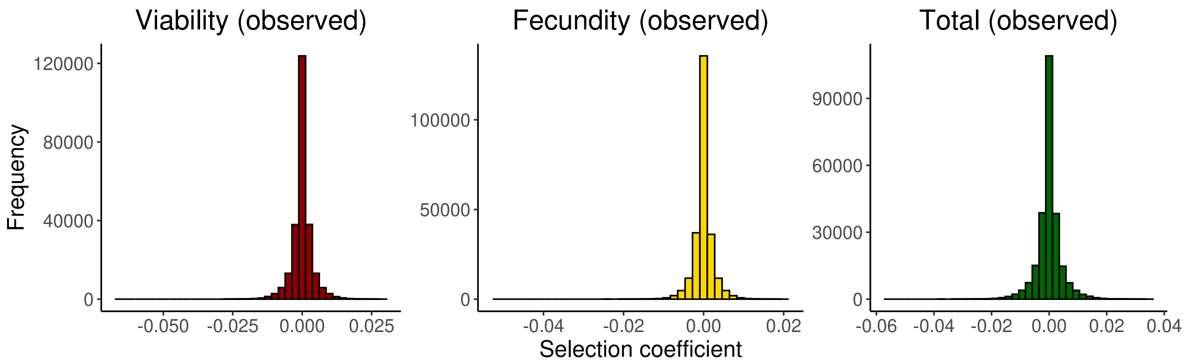


**Figure B.2.1:** Observed distributions of $\hat{s}$ for viability (dark red), fecundity (yellow), and total (dark green) selection.

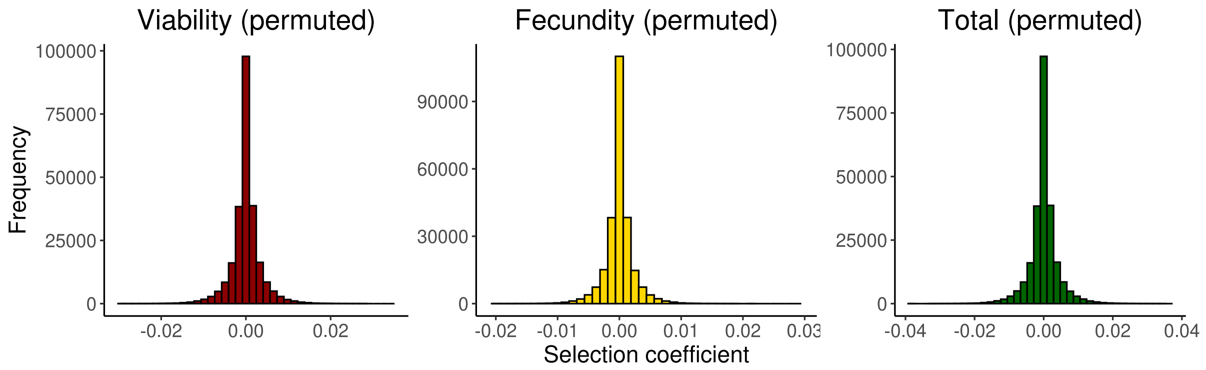


**Figure B.2.2:** Sex-label permuted distributions of $\hat{s}$ for viability (dark red), fecundity (yellow), and total (dark green) selection.

#### B.3. Genome-wide bootstrapped $p$ values for SDS

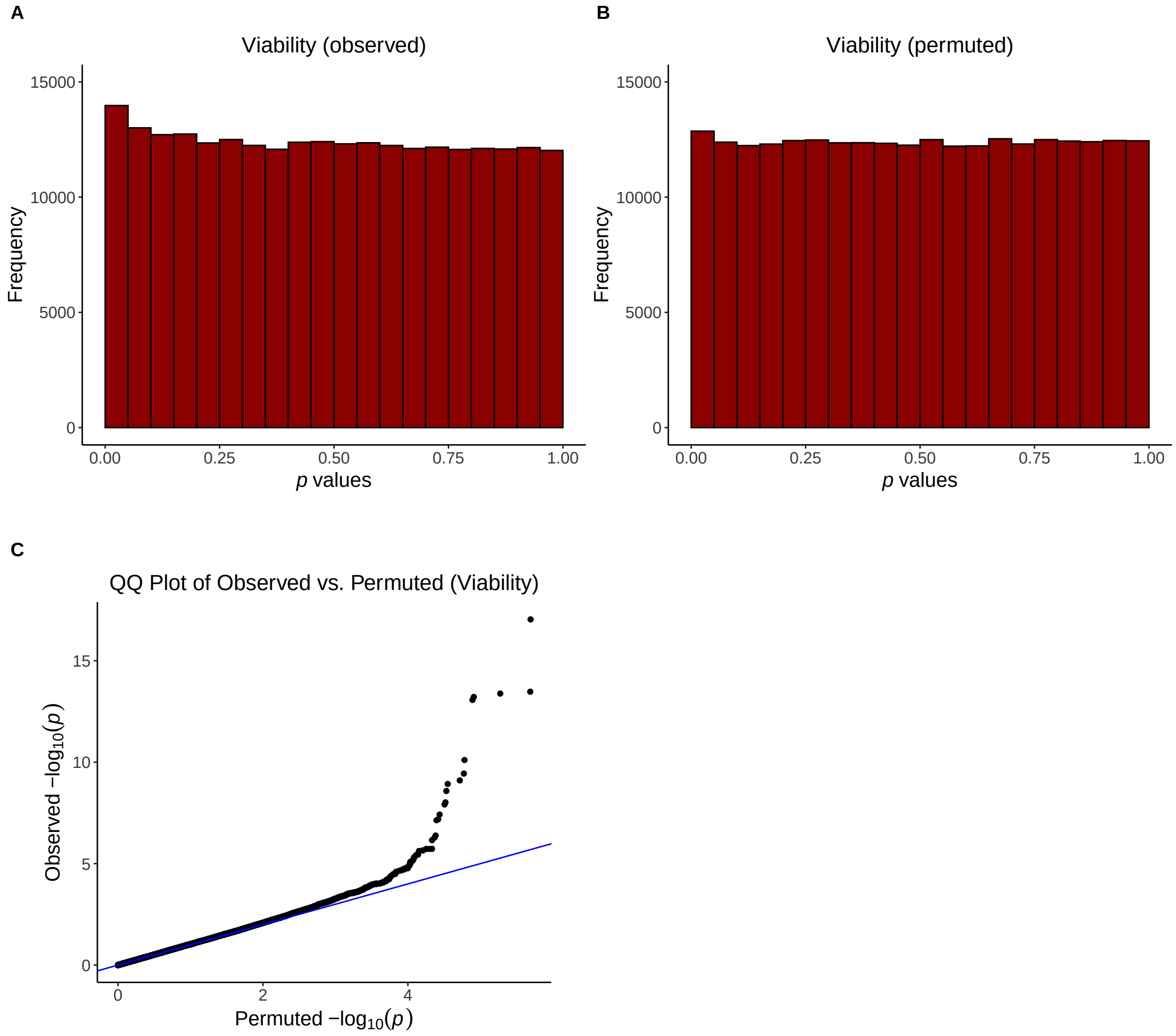


**Figure B.3.1:** Bootstrap $p$ value distributions for the observed (A) and permuted (B) $\hat{s}$ values for viability selection. The Quantile-Quantile (QQ) plot (C) compares the observed vs permuted bootstrapped $p$ values, -log transformed.

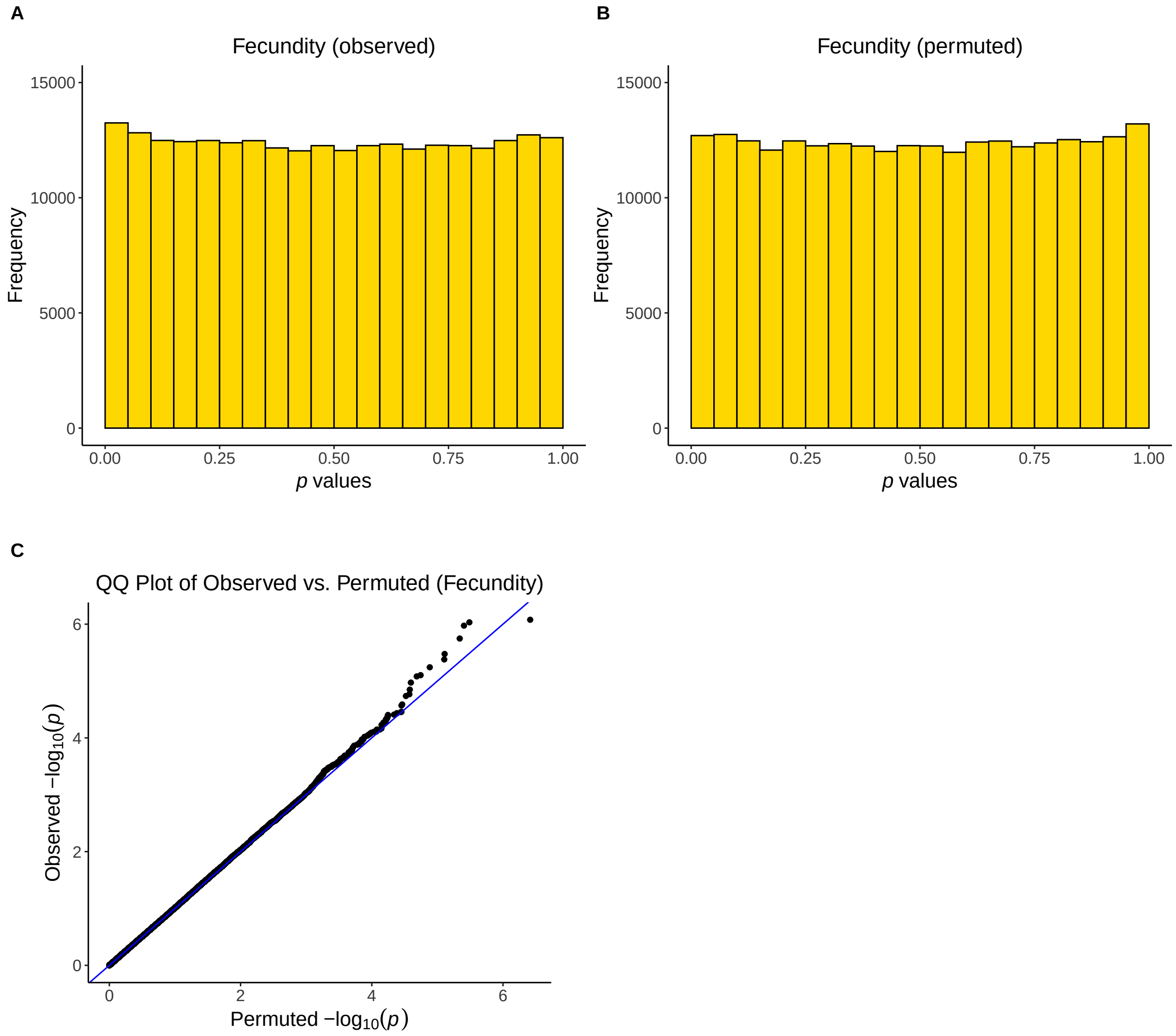


**Figure B.3.2:** Bootstrap $p$ value distributions for the observed (A) and permuted (B) $\hat{s}$ values for fecundity selection. The Quantile-Quantile (QQ) plot (C) compares the observed vs permuted bootstrapped $p$ values, -log transformed.

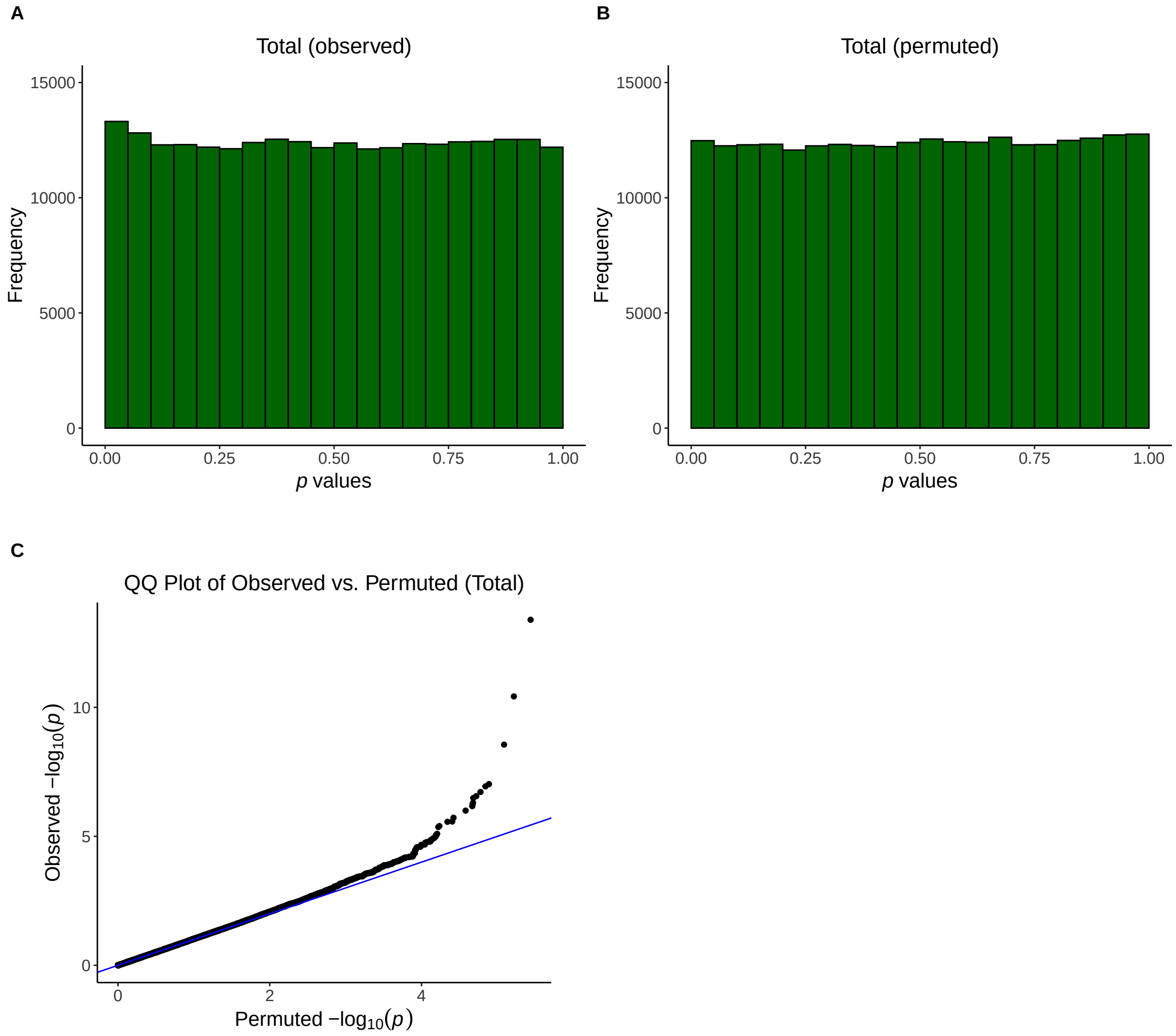


**Figure B.3.3:** Bootstrap $p$ value distributions for the observed (A) and permuted (B) $\hat{s}$ values for total selection. The Quantile-Quantile (QQ) plot (C) compares the observed vs permuted bootstrapped $p$ values, -log transformed.

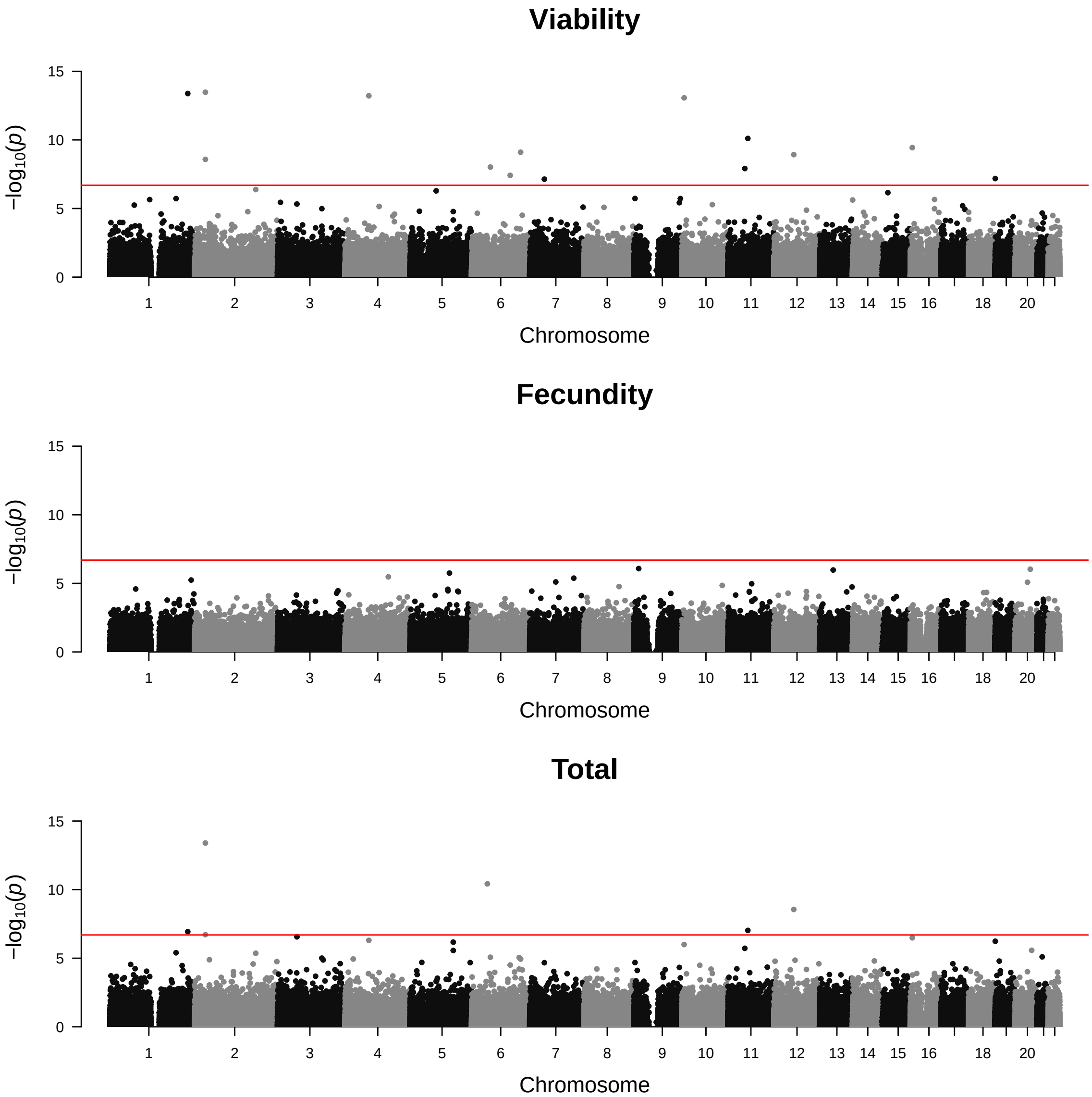


**Figure B.3.4:** Manhattan plots for the observed $p$ values for viability (top), fecundity (middle), and total (bottom) selection. The -log$_{10}$-transformed $p$ values are displayed on the y-axis for windows on each autosome. The red line indicates the Bonferroni threshold ($p = 2 \times 10^{-7}$).

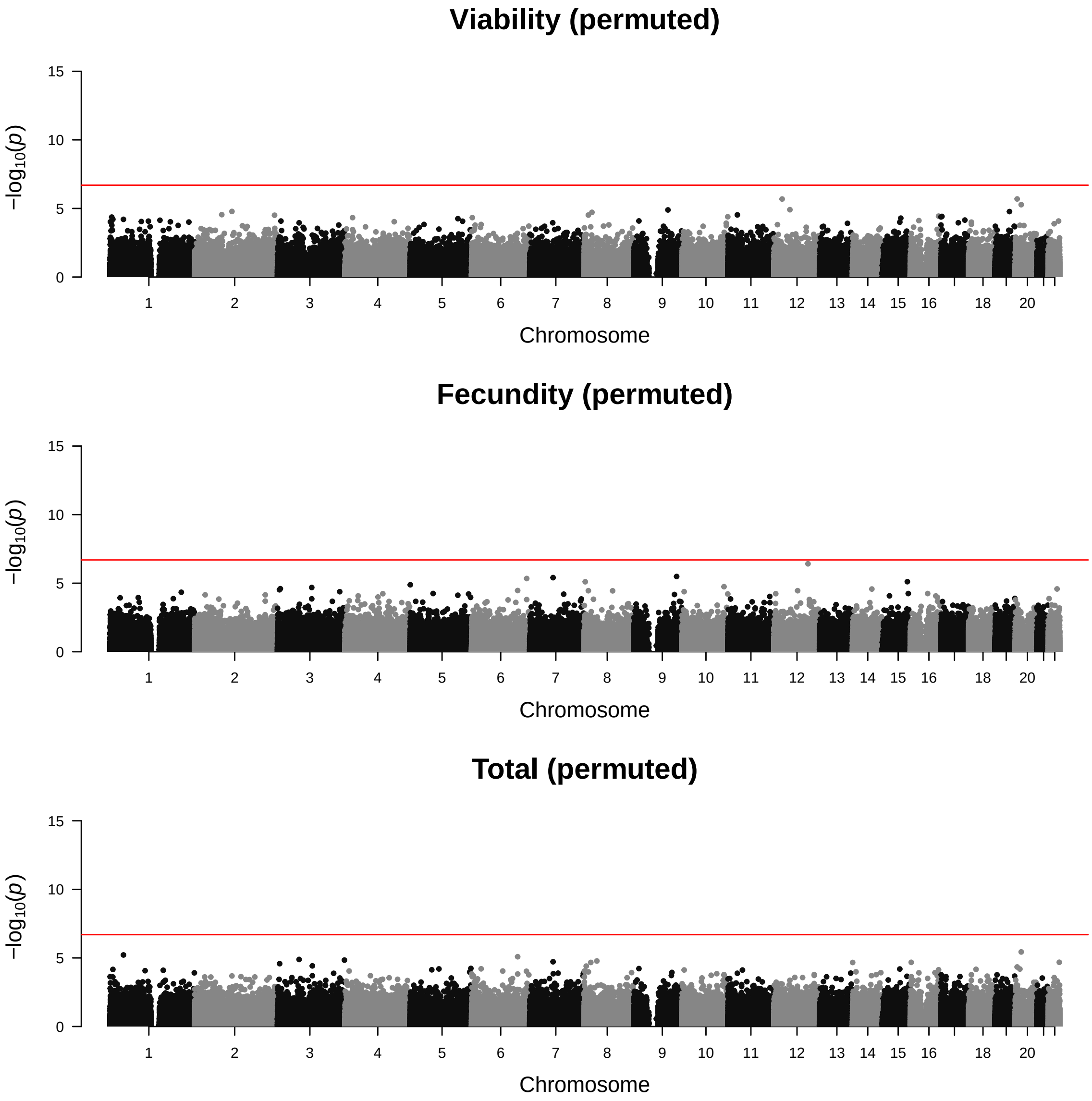


**Figure B.3.5:** Manhattan plots for the permuted $p$ values for viability (top), fecundity (middle), and total (bottom) selection. The -$\log_{10}$-transformed $p$ values are displayed on the y-axis for windows on each autosome. The red line indicates the Bonferroni threshold ($p = 2 \times 10^{-7}$).

#### 335 B.4. Functional consequences of windows significant for SDS

**Table B.4.1:** The 15 significant windows after non-sequential Bonferroni correction (at $\alpha = 0.05$) for viability and their accompanying flanking SNPs. SNPs were annotated using Ensembl's Variant Effect Predictor (VEP) to retrieve associated genes and phenotypes.

| Window | P value (Bonferroni) | SNP | MAF | Location | Associated Gene(s) | Associated Phenotypes |
|---|---|---|---|---|---|---|
| 16851 | 2.27307E-12 | rs9357509 | 0.44 | 6:46485635 | RP11-795J1.1 | - |
| | 2.27307E-12 | rs74932938 | 0.02 | 6:46486527 | RP11-795J1.1 | - |
| 7343 | 8.34305E-09 | rs9679162 | 0.4 | 2:31247514 | GALNT14 | Various cancers[19] |
| | 8.34305E-09 | rs5009910 | 0.4 | 2:31249014 | GALNT14 | Various cancers[19] |
| 38563 | 1.02819E-08 | rs11586639 | 0.45 | 1:228715705 | - | - |
| | 1.02819E-08 | rs10916343 | 0.21 | 1:228727961 | - | - |
| 13209 | 1.50915E-08 | rs62306016 | 0.13 | 4:70243367 | - | - |
| | 1.50915E-08 | rs12500647 | 0.13 | 4:70261469 | RP11-790I12.1 | - |
| 2195 | 2.11826E-08 | rs17399746 | 0.02 | 6:6876020 | LINC00707 | - |
| | 2.11826E-08 | rs10905035 | 0.18 | 6:6883788 | LINC00707 | - |
| 11696 | 1.94498E-05 | rs35211059 | 0.2 | 11:58542727 | - | - |
| | 1.94498E-05 | rs11605651 | 0.49 | 11:58546907 | - | - |
| 1883 | 9.04902E-05 | rs72772869 | 0.07 | 16:6134402 | RBFOX1, RP11-509E10.1, RP11-420N3.2 | RBFOX1-related neurodevelopmental |
| | 9.04902E-05 | rs889700 | 0.43 | 16:6134549 | RBFOX1, RP11-509E10.1, RP11-420N3.2 | RBFOX1-related neurodevelopmental disorder |
| 32015 | 0.000196351 | rs74957359 | 0.03 | 6:144034840 | PHACTR2 | Transient neonatal diabetes mellitus |
| | 0.000196351 | rs1015339 | 0.38 | 6:144034848 | PHACTR2 | Transient neonatal diabetes mellitus |
| 11261 | 0.000293234 | rs4403838 | 0.34 | 12:58439028 | - | - |
| | 0.000293234 | rs1506885 | 0.3 | 12:58440197 | - | - |
| 7342 | 0.000651206 | rs10209881 | 0.4 | 2:31246249 | GALNT14 | Various cancers[19] |
| | 0.000651206 | rs9679162 | 0.4 | 2:31247514 | GALNT14 | Various cancers[19] |
| 18389 | 0.00237479 | rs9475319 | 0.49 | 6:55361878 | HMGCLL1 | - |
| | 0.00237479 | rs12210883 | 0.29 | 6:55366664 | HMGCLL1 | - |
| 10905 | 0.003012316 | rs35341123 | 0.04 | 11:49554625 | - | - |
| | 0.003012316 | rs1814175 | 0.43 | 11:49559172 | - | Height |
| 26815 | 0.0095051 | rs34118295 | 0.03 | 6:113526446 | - | |
| | 0.0095051 | rs4945931 | 0.34 | 6:113528108 | - | |
| 174 | 0.016250667 | rs62132585 | 0.19 | 19:561773 | - | - |
| | 0.016250667 | rs62132587 | 0.29 | 19:561816 | - | - |
| 10100 | 0.018030531 | rs7432 | 0.06 | 7:42977342 | MRPL32 | - |
| | 0.018030531 | rs655367 | 0.23 | 7:42991064 | MRPL32, AC005537.2 | - |

#### B.5. Comparison of SDS at single sites to Ruzicka *et al.*

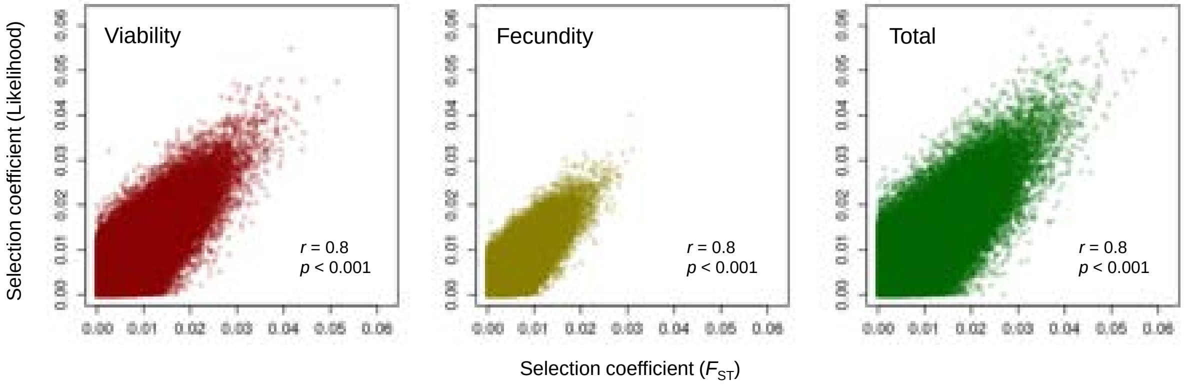


**Figure B.5.1:** Shown are scatter plots comparing our maximum likelihood estimates of the selection coefficients (using estimators in equations (SI 6) and (SI 16) at single sites (y-axis) to selection coefficients derived from Ruzicka *et al.* (2)'s $F_{\mathrm{ST}}$ estimates (see equation (SI 17)) on the x-axis. Estimates are from genotyped array SNPs that overlapped our datasets (190,374 SNPs, $N = 303,824$ in our study, $N = 249,021$ in Ruzicka *et al*). The Pearson's correlation coefficients ($r$) and their respective $p$ values are shown for each of the three modes of selection: viability (dark red), fecundity ("reproductive $F_{\mathrm{ST}}$"; gold), and total ("gametic $F_{\mathrm{ST}}$"; dark green). Comparisons indicate that our estimators are broadly consistent.

#### B.6. Sexually antagonistic selection on fecundity at single sites

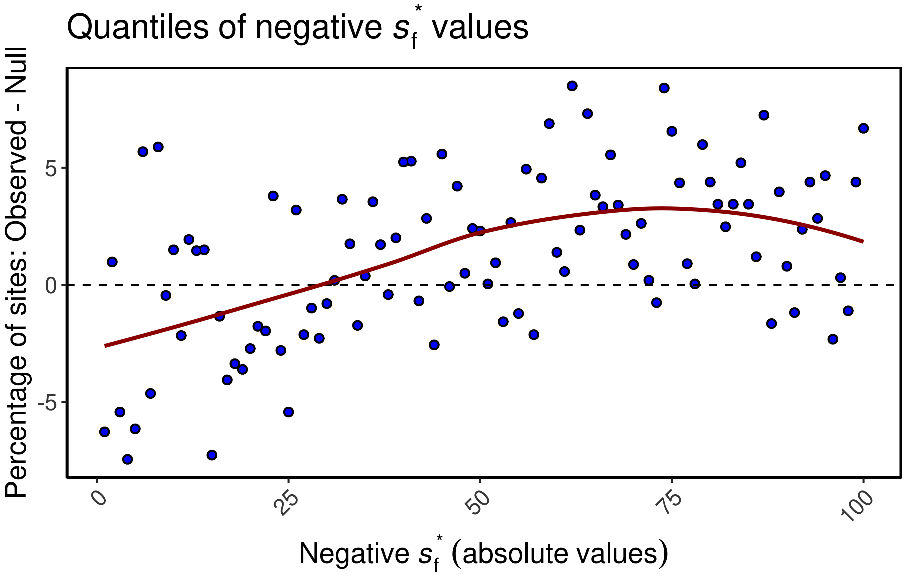


**Figure B.6.1:** The excess of individual sites showing evidence for sexually antagonistic selection relative to the empirical null. The absolute values of all negative $s_f^*$ (see equation (SI 19)) from the empirical null are binned into 100 quantiles, shown on the x-axis. As with the haplotype analysis, empirical nulls were generated by permuting the sex labels. The y-axis shows the percentage excess or deficit of $s_f^*$ in each bin from the observed data. A LOESS curve is added for emphasis.

#### B.7. Weighted standard major axis regressions on physiological traits

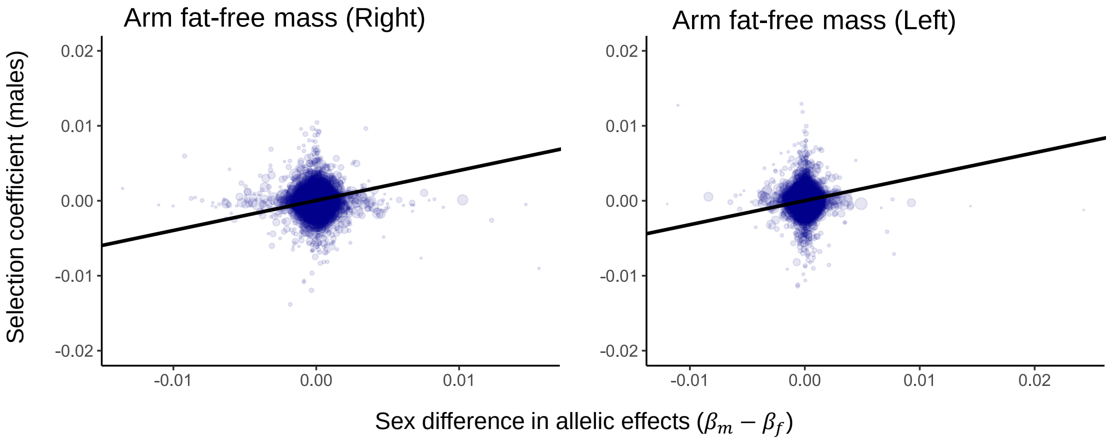


**Figure B.7.1:** Shown are two examples of the weighted standard major axis regression used to estimate the strength of sexually antagonistic selection on alleles associated with 27 physiological traits. The y-axis represents the estimated selection coefficient in males, whereas the x-axis shows the sex difference in variant effects on the trait. Each point represents a SNP and the size is proportional to the regression weighting given by equation (SI 26). In the above two cases, the selection coefficients are estimated on fecundity.

#### B.8. Estimating the strength and frequency of SDS using ABC

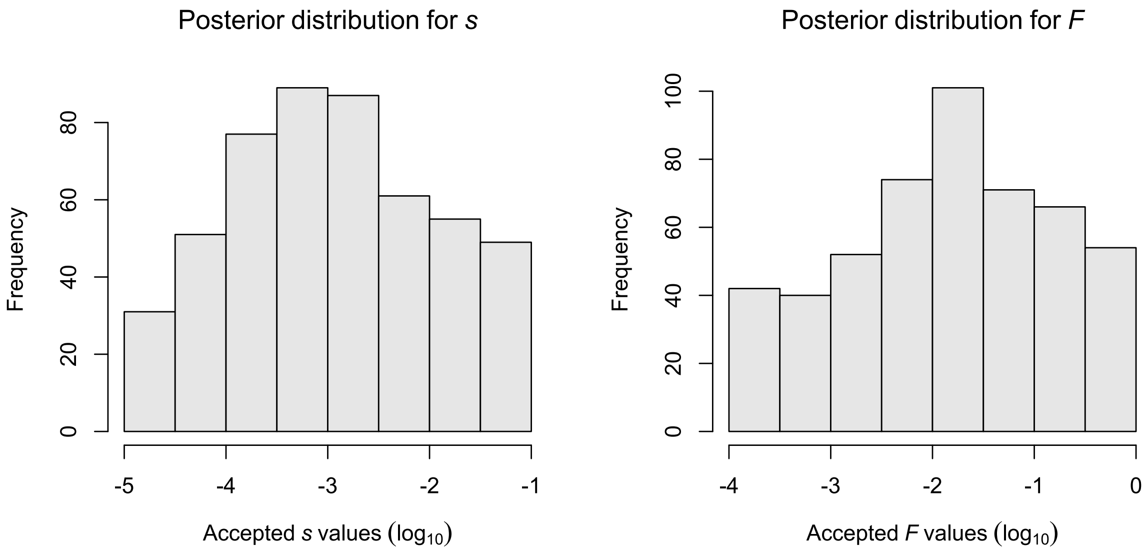


**Figure B.8.1:** The marginal posterior distributions for the selection coefficient $s$ (left) and the frequency of selection $F$ (right) obtained using Approximate Bayesian Computation. Each distribution shows the 500 values retained at an acceptance threshold of 1% after 50,000 runs.

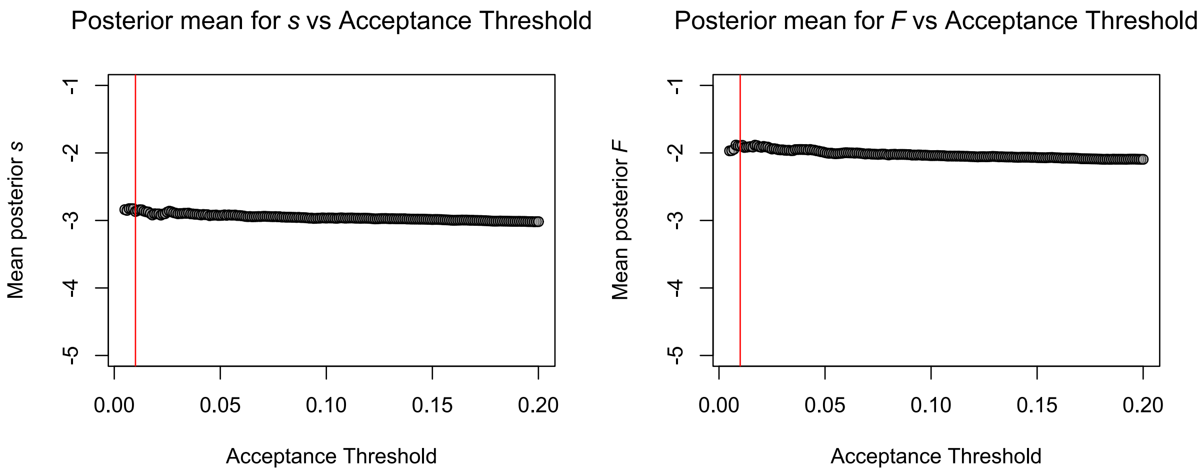


**Figure B.8.2:** The mean of the marginal posterior distributions for $s$ (left) and $F$ (right) at varying acceptance thresholds after 50,000 runs. The red line shows an acceptance threshold of 1% as used in the main analysis. Mean posterior estimates are relatively robust to the number of accepted posterior values.

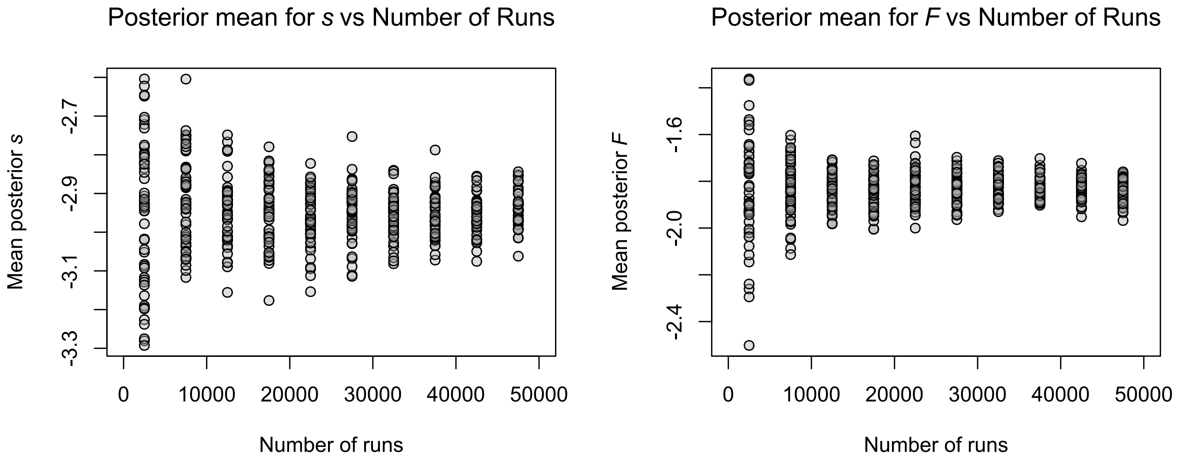


**Figure B.8.3:** The mean of the marginal posterior distributions for $s$ (left) and $F$ (right) with varying numbers of simulated prior values. Values were obtained by downsampling from 50,000 runs with replacement. This was done 50 times for each of 10 sample sizes, ranging from 5% to 95% of the total number of runs. This resulted in 50 means for each downsample percentage. Marginal posterior mean estimates are relatively insensitive to the number of runs.
